# Doublecortin and JIP3 are neural-specific counteracting regulators of dynein-mediated retrograde trafficking

**DOI:** 10.1101/2022.08.10.503449

**Authors:** Lu Rao, Peijun Li, Xinglei Liu, Qi Wang, Alexander I. Son, Arne Gennerich, Judy S. Liu, Xiaoqin Fu

**Author notes:** These authors contributed equally (Lu Rao, Peijun Li).

## Abstract

Mutations in the microtubule (MT)-binding protein doublecortin (DCX) or in the MT-based molecular motor dynein result in lissencephaly. However, a functional link between DCX and dynein has not been defined. Here, we demonstrate that DCX negatively regulates dynein-mediated retrograde transport by reducing dynein’s association with MTs and by disrupting the composition of the dynein motor complex. Previous work showed an increased binding of the adaptor protein C-Jun-amino-terminal kinase-interacting protein 3 (JIP3) to dynein in the absence of DCX. Using purified components, we demonstrate that JIP3 forms an active motor complex with dynein and its cofactor dynactin with two dyneins per complex. DCX competes with the binding of the second dynein, resulting in a velocity reduction of the complex. We conclude that DCX negatively regulates dynein-mediated retrograde transport through two critical interactions by regulating dynein binding to MTs and by regulating the association of JIP3 to the dynein motor complex.

## INTRODUCTION

Lissencephaly is a cortical malformation characterized by a “smooth cortex” which arises from disruption of normal development (des Portes et al., 1998; Deuel et al., 2006; Francis et al., 1999; Gleeson et al., 1998). Patients with lissencephaly often have an associated microcephaly indicating defects in neural progenitor proliferation (Lee et al., 2010; Pramparo et al., 2010). Furthermore, the “smooth” brain is the result of abnormal neuronal migration, which causes an abnormally thick 4-layered cortex (Dobyns et al., 1984; Jellinger and Rett, 1976), and many patients with lissencephaly also have a reduction or absence of major axon tracts indicating problems with axon guidance or outgrowth (Kappeler et al., 2007). Thus, causative genes for lissencephaly encode proteins with critical roles in each of these steps in development: neural progenitor proliferation, neuronal migration, and axon outgrowth. Defining the molecular and cellular functions of lissencephaly genes are therefore critical for understanding early human neural development.

Interestingly, many of the causative genes for lissencephaly encode proteins related to the microtubule (MT) cytoskeleton. These include doublecortin (DCX), a MT-binding protein (des Portes et al., 1998; Gleeson et al., 1998); tubulin α1a, a major subunit of MTs (Keays et al., 2007); cytoplasmic dynein (hereinafter, “dynein”) heavy chain (DHC), the main retrograde motor protein (Poirier et al., 2013); and the dynein co-factor, lissencephaly1 (Lis1) (Reiner et al., 1993). This suggests that these MT proteins may be functionally related during neuronal development. Indeed, DCX overexpression can rescue nucleus–centrosome coupling defect and neuronal migration defect caused by the disruption of dynein/Lis1 function in mouse cerebellar granule neurons (Tanaka et al., 2004). However, the molecular mechanisms through which these MT-related proteins are functional related are only partly understood.

Both DCX and doublecortin-like kinase 1 (DCLK1) regulate MT-based motor transport mediated by the Kinesin-3 family motor KIF1A (Deuel et al., 2006; Lipka et al., 2016; Liu et al., 2012). As DCX binds the surface of the MT lattice (Bechstedt and Brouhard, 2012; Fourniol et al., 2010), it is logical to hypothesize that DCX regulates axonal transport by modifying the interactions of molecular motors with MTs as they step along the MT lattice. DCX was found to be part of the dynein motor complex (Li et al., 2021; Tanaka et al., 2004) and influence the association between dynein and c-Jun NH2- terminal kinase (JNK)-interacting protein-3 (JIP3), an adaptor protein of kinesin and dynein that mediates both anterograde and retrograde transport (Arimoto et al., 2011; Drerup and Nechiporuk, 2013; Li et al., 2021), implying that DCX may influence other aspects of dynein function.

In the absence of MTs, dynein assumes an auto-inhibited “inverted” conformation (Torisawa et al., 2014; Toropova et al., 2017) and upon binding to its largest co-factor dynactin together with a cargo adaptor such as Bicaudal D2 (BicD2), converts into a parallel conformation capable of binding MTs (Chowdhury et al., 2015; McKenney et al., 2014; Olenick et al., 2016; Schlager et al., 2014; Zhang et al., 2017). Dynein-dynactin- BicD2 (DDB) complex formation is facilitated by Lis1, which interacts with dynein’s motor domain and prevents its auto-inhibitory conformation (Marzo et al., 2020). However, whether dynein and dynactin can form an active motor complex with JIP3, remains unknown.

In this study, we show that DCX plays critical roles in dynein-mediated retrograde transport in axons through two different mechanisms: First, DCX decreases dynein binding to MTs, and second, DCX regulates the association of dynein with JIP3. We further demonstrate for the first time the formation of ultra-processive dynein-dynactin-JIP3 (DDJ) motor complexes with up to two dyneins and show that DCX displaces the second dynein from a DDJ complex, resulting in a reduction of the velocity of the DDJ motor complex. Together, we demonstrate that DCX plays key roles in axon-based transport to mediate the highly specific trafficking of proteins in both anterograde and retrograde directions during neuronal growth and development by modulating the activity of MT-plus and minus-end-directed motor proteins.

## RESULTS

### Dynein-mediated retrograde trafficking increases in the absence of DCX

Our previous work showed that DCX is essential for the function of the MT plus-end-directed kinesin-3 motor KIF1A and regulates its anterograde trafficking (Liu et al., 2012). DCX has also been shown to associate with the MT minus-end-directed motor dynein (Tanaka et al., 2004). In addition, we recently reported that dynein is involved in the DCX-mediated trafficking of Golgi extensions into dendrites, suggesting a functional link between DCX and dynein (Li et al., 2021). To determine whether functional interactions between DCX and dynein exist, we tested whether DCX affects the retrograde trafficking of dynein.

To visualize dynein function *in vivo*, we first transfected either WT, *Dcx-/y*; or *Dcx-/y;Dclk-/-* dissociated cortical neuronal cultures with a construct expressing RFP-tagged, neuron-specific dynein intermediate chain (DIC) isoform IC-1B on DIV (days *in vitro*) 6 (Ha et al., 2008). Time-lapse imaging of DIC IC-1B-RFP was performed on DIV8 to visualize dynein motor activity directly in axons. Recorded images were converted to kymographs (Fig. 1A). For all calculations and measurements of dynein-mediated movement, DIC above an intensity threshold located in the proximal region of axons (∼ 100 µm away from cell body) were analyzed. A complex was counted as mobile only if the displacement was at least 5 µm over the course of the 180 seconds, otherwise it was counted as stationary. Distribution calculations of DIC mobility status (anterograde, retrograde and stationary) demonstrate that mobile DIC predominantly display retrograde movements in axons, and the percentages or run frequency of moving dynein complexes are similar in different neurons (Fig. 1B). Remarkably, both the run length (the average distance traveled during the recorded time period) and the velocity of the fluorescently-tagged dynein complexes were significantly increased in both *Dcx-/y* and *Dcx-/y;Dclk1-/-* axons compared to WT axons (Fig. 1C, Suppl. Videos 1-2) (run length and velocity distributions of retrograde moving DIC in different neurons are shown in Suppl. Fig. 1A). Reintroduction of DCX fully rescued the retrograde trafficking of DIC observed in *Dcx-/y* neurons (Fig. 1C, P is 0.57 and 0.18 for DCX rescue compared to WT for DIC run length and speed, respectively). In contrast, DCLK1, a doublecortin domain-containing protein that is structurally similar to DCX, only partially rescued the phenotype (Fig. 1C, P is 0.07 and 0.25 for DCLK1 rescue compared to Dcx-/y for DIC run length and speed, respectively).

**Figure 1.**
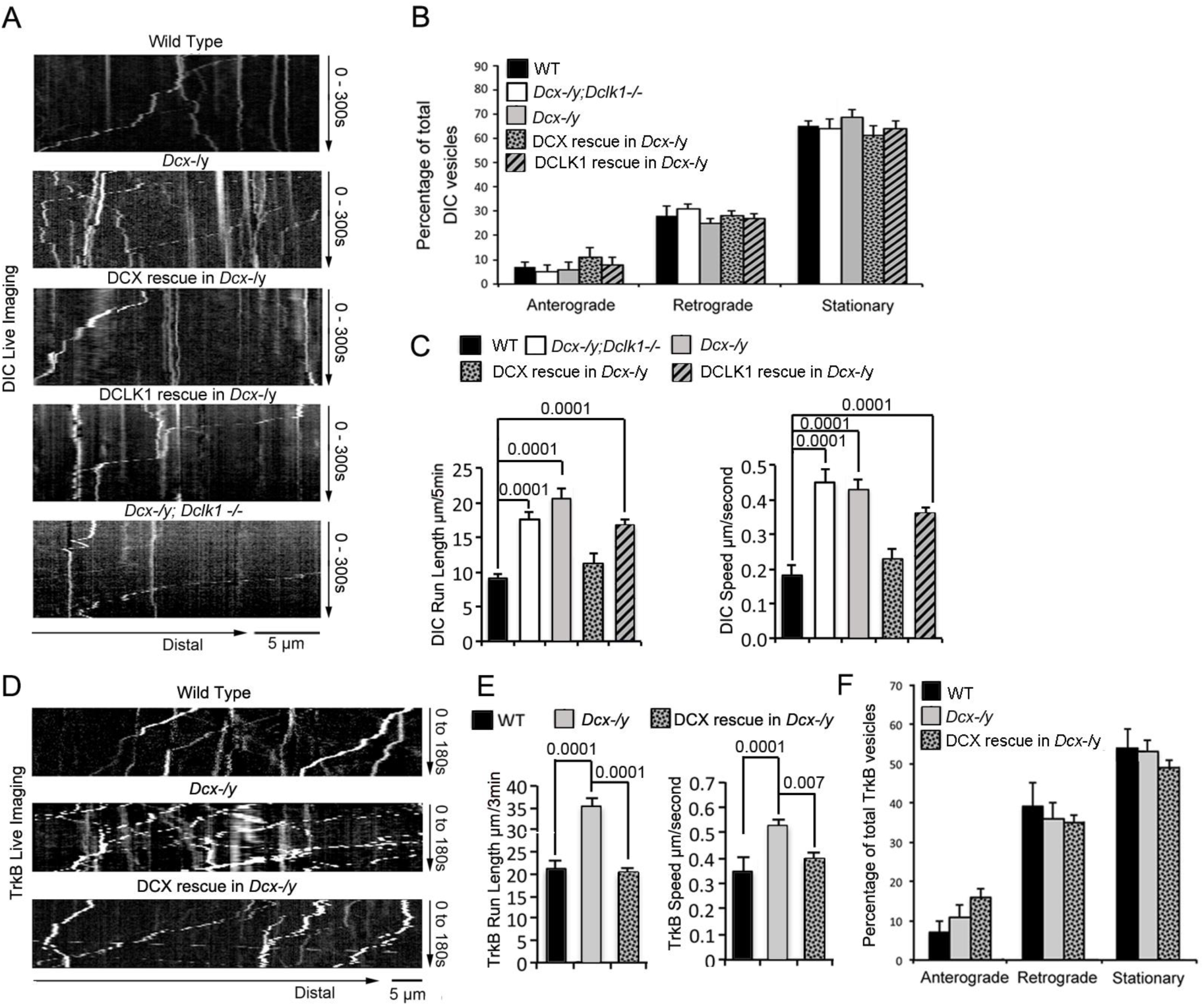
The retrograde trafficking of the dynein motor and TrkB transport is increased in axons without DCX. (A) WT, Dcx-/y or Dcx-/y;Dclk1-/- associated cortical neuronal culture were transfected with plasmids expressing DIC-RFP on DIV6 and imaged on DIV8. For rescue experiments, Dcx-/y neurons were transfected with plasmids expressing DIC-RFP combined with plasmids expressing either DCX-GFP or DCLK1-GFP. Representative kymographs of DIC-RFP transport in axons are shown. (B) Distribution calculations of the DIC vesicle mobility status (anterograde, retrograde and stationary) are demonstrated. No significant differences are observed among different neurons. (C) Quantifications of DIC-RFP run length within 300 seconds and velocity are shown. DCX, but not DCLK1, fully rescued the increased dynein motor transport observed in DCX deficient axons. P values comparing WT and DCX rescue for run length and speed are 0.57 and 0.18, respectively. Other P values are shown in the figure. (D) Dissociated cortical neuronal cultures from WT or Dcx-/y mice were transfected with plasmids expressing TrkB-RFP with/without plasmids expressing DCX-GFP on DIV6 and imaged on DIV8. Representative kymographs of TrkB-RFP transport in axons are shown. (E) Quantification of vesicle Run length within 180 seconds and velocity are demonstrated. DCX rescued the increased TrkB-RFP transport in DCX deficient axons. (F) Distribution calculations of the TrkB vesicle mobility status (anterograde, retrograde and stationary) are demonstrated. No significant differences are observed among different neurons. Data are based on three independent experiments of each condition. P-values from t-tests are shown in each panel. Total numbers of neurons (N) and vesicles (V) used in the calculations are indicated in Suppl. Fig. 1. See also Suppl. Videos 1-4.

To determine whether the dynein-motility changes we see in neurons can also be observed for a physiologically relevant dynein cargo, we tested the retrograde trafficking of Tropomyosin receptor kinase B (TrkB), the neurotrophin receptor whose retrograde transport is mediated by dynein (Ha et al., 2008; Heerssen et al., 2004; Yano et al., 2001; Zhou et al., 2012). As with IC-1B, the run length and velocity of retrogradely moving TrkB are also significantly increased in *Dcx-/y* axons (Fig. 1D and 1E, Suppl. Videos 3-4), and reintroduction of DCX into *Dcx-/y* neurons rescued the phenotype (Fig. 1E, P is 0.64 and 0.057 for DCX rescue compared to WT for TrkB run length and speed, respectively). Like IC-1B, the majority of mobile TrkB vesicles in axons were transported in retrograde direction (Fig. 1F) and no significant differences were found between the percentage of vesicles measured under WT, *Dcx-/y*, and rescue conditions for anterograde and retrograde moving particles and for stationary vesicles (Fig. 1F) (run length and velocity distributions of IC-1IB and TrkB vesicles are shown in Suppl. Fig. 1B). Overall, our data indicate that loss of DCX increases dynein-mediated vesicular retrograde transport in the axon.

### The effect of DCX on retrograde transport is mediated through direct interactions between DCX and dynein

In addition to its binding to MTs, DCX also associates with the dynein motor complex (Tanaka et al., 2004; Taylor et al., 2000). Pull down assay confirms there exists direct interaction between DCX and cytoplasmic dynein intermediate chain 2 (Table 1). We therefore hypothesized that DCX influences dynein movement through direct interactions. To confirm that DCX interacts with dynein and to define which domain of DCX is critical for its association with dynein, we expressed HA-tagged full-length DCX (FL-DCX), an N-terminal DCX construct (N-DCX) containing the R1 and R2 MT-binding domains (amino acids 1-270) or a C-terminal construct containing the Serine/Proline (SP) rich domain of DCX (C-DCX) (amino acids 271-361, Fig. 2A) in HEK293 cells. Consistent with the previous report (Tanaka et al., 2004), DIC precipitated with FL-DCX (Fig. 2B). Interestingly, more DIC precipitated with N-DCX than FL-DCX (Fig. 2B); similarly, N- DCX showed an increased immunoprecipitation of the DHC compared to FL-DCX (Suppl. Fig. 2A). This result suggests that N-DCX has a stronger affinity for the dynein motor complex than FL-DCX. We reasoned that if the interaction of DCX with dynein plays an important role in regulating dynein function, then N-DCX should have a stronger effect on regulating dynein-mediated retrograde transport than FL-DCX. Indeed, introducing N- DCX either into DCX knockout neurons (Fig. 2C-D and Suppl. Fig. 3A-B) or WT neurons (Fig. 2F-G and Suppl. Fig. 3C-D) decreases the retrograde transport of TrkB to a greater extent than FL-DCX. Our results suggest that DCX decreases dynein-mediated retrograde transport through direct interactions with dynein motor complex.

**Figure 2.**
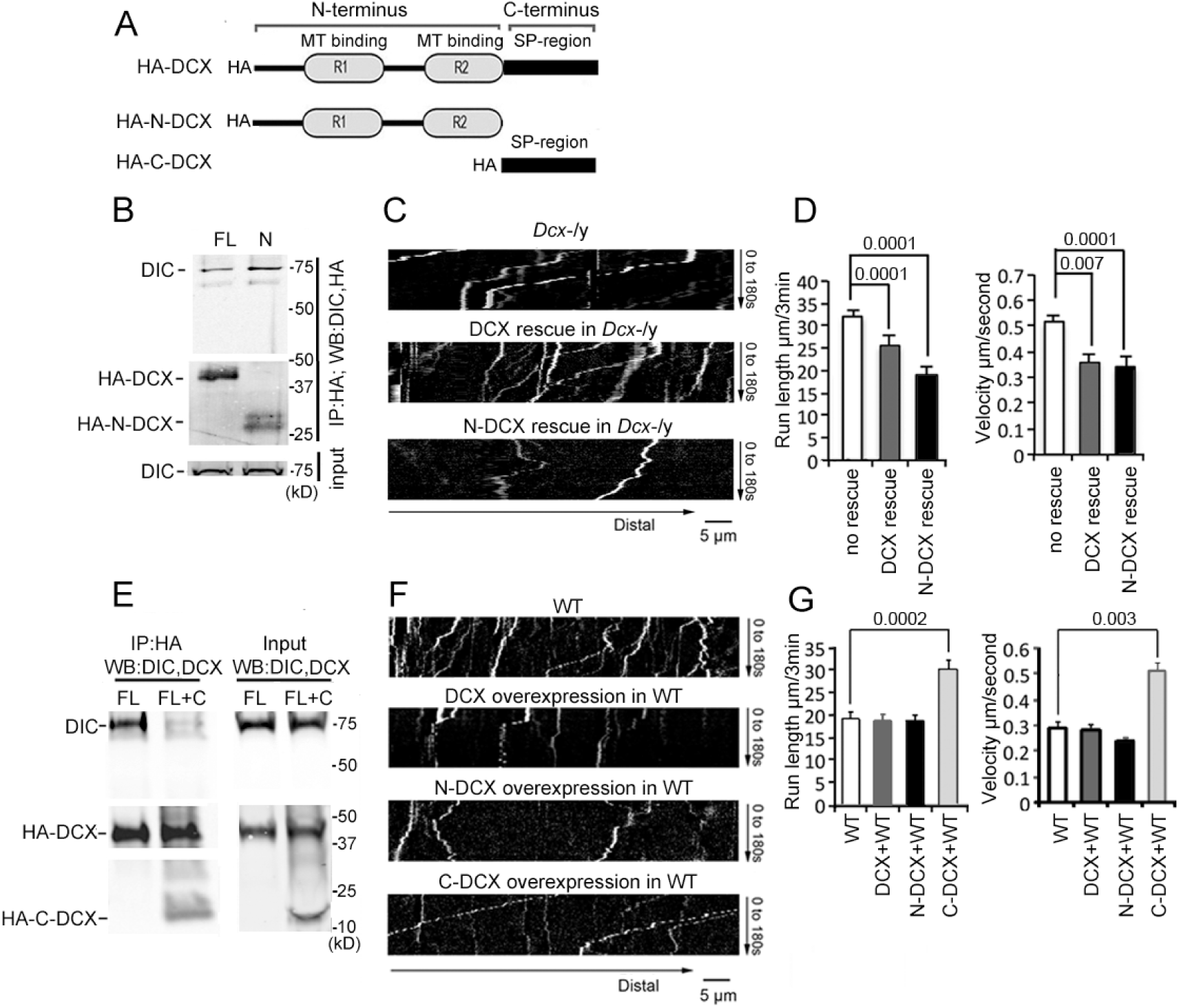
DCX affects the retrograde transport through DCX/dynein interaction. (A) A schematic of DCX protein domain structure. N-DCX has N terminal R1 and R2 domains represent MT-binding domains of DCX, while C-DCX has DCX C terminal Serine/Proline (SP) rich domain. (B) More N-DCX proteins are pulled down with DIC compared to full length DCX. HEK cells were transfected with plasmids expressing HA tagged either full length DCX (FL) or N-DCX (N) for two days. Protein lysates were used for immunoprecipitation using antibodies for HA and analyzed by Western blot for DIC and HA. (C) Dissociated cortical neuronal culture from Dcx-/y;Dclk1-/- mice were transfected with plasmids expressing TrkB-RFP with/without plasmids expressing DCX or N-DCX on DIV6 and imaged on DIV8. Representative kymographs of TrkB-RFP transports in axons are shown. (D) The expression of either full length DCX or N-DCX in DCX knockout neurons significantly decreased TrkB retrograde transport while N-DCX has stronger effect compared to full length DCX. (E) C-DCX decreases DCX/DIC association. HEK cells were transfected with plasmids expressing HA tagged full length DCX (FL) with/without plasmid expressing C-DCX for two days. Protein lysates were used for immunoprecipitation using antibodies for HA and analyzed by Western blot for DIC and HA. (E) Dissociated cortical neuronal culture from wild type mice were transfected with plasmids expressing TrkB-RFP with/without plasmids expressing DCX, N-DCX or C- DCX on DIV6 and imaged on DIV8. Representative kymographs of TrkB-RFP transports in axons are shown. (F) Run length within 180 seconds and velocity distributions of retrograde TrkB complexes in axons are quantified. C-DCX overexpression in wild type neurons mimicked the phenotype of TrkB retrograde trafficking observed in Dcx-/y axons. All quantification data is based on three independent experiments of each condition. P-values from t-tests are shown in each panel. Total numbers of neurons (N) and vesicles (V) used in the calculations are indicated in Suppl. Fig. 3. See also Suppl. Fig. 2.

**Table 1:** Pull down assay results show cytoskeleton proteins associated with DCX.

| Protein IDs | Protein names | Q-value | Score | Average normalized intensity |  |
| --- | --- | --- | --- | --- | --- |
|  |  |  |  | Control (HA) | HA-DCX |
| Q61301 | Catenin alpha-2 | 0 | 83 | 0 | 158960000 |
| P57780 | Alpha-actinin-4 | 0 | 76 | 0 | 246655000 |
| Q8BMK4 | Cytoskeleton-associated protein 4 | 0 | 70 | 0 | 124115000 |
| Q02248 | Catenin beta-1 | 0 | 55 | 0 | 133995000 |
| P05213 | Tubulin alpha-1B chain;Tubulin alpha-4A chain | 0 | 52 | 0 | 600270000 |
| Q9D6F9 | Tubulin beta-4A chain | 0 | 38 | 0 | 91708000 |
| Q8BTM8 | Filamin-A | 0 | 38 | 0 | 48710000 |
| Q8K341 | Alpha-tubulin N-acetyltransferase 1 | 0 | 36 | 0 | 9883000 |
| Q99KJ8 | Dynactin subunit 2 | 0 | 32 | 0 | 67364000 |
| Q3TPJ8 | Cytoplasmic dynein 1 intermediate chain 2 | 0 | 25 | 0 | 34566000 |
| Q9CPW4 | Actin-related protein 2/3 complex subunit 5 | 0 | 21 | 0 | 32742000 |
| Q9D898 | Actin-related protein 2/3 complex subunit 5-like protein | 0 | 17 | 0 | 3859300 |
| Q6R891 | Neurabin-2 | 0 | 15 | 0 | 6523000 |
| P28667 | MARCKS-related protein | 0 | 14 | 0 | 52599500 |
| Q9JM76 | Actin-related protein 2/3 complex subunit 3 | 0 | 14 | 0 | 30507500 |
| Q7TPR4 | Alpha-actinin-1 | 0 | 12 | 0 | 6880500 |
| P60710 | Actin, cytoplasmic 1, N-terminally processed | 0 | 9 | 0 | 30897500 |
| Q3UX10 | Tubulin alpha chain-like 3 | 0 | 8 | 0 | 7851000 |
| Q9CQV6 | Microtubule-associated proteins 1A/1B light chain 3B | 0 | 7.5 | 0 | 42111500 |
| Q922F4 | Tubulin beta-6 chain | 0.001 | 7.3 | 0 | 12908500 |
| Q9QZB9 | Dynactin subunit 5 | 0.007 | 6.2 | 0 | 2061750 |

### The C-terminal S/P-rich domain of DCX decreases DCX-dynein interactions

In contrast to FL-DCX and N-DCX, C-DCX did not immunoprecipitate with dynein (Suppl. Fig. 2B-C), consistent with previous report (Taylor et al., 2000). Since N- DCX, which misses the C-terminal domain, has a stronger affinity for dynein than FL- DCX, we hypothesized that the C-terminus of DCX inhibits the interaction between DCX and dynein. Indeed, in the presence of C-DCX, significantly less DIC precipitated with FL- DCX (Fig. 2E); similarly, less DHC was precipitated with either FL-DCX or N-DCX in the presence of C-DCX (Suppl. Fig. 3B-C). Furthermore, C-DCX overexpression in WT neurons significantly increased dynein-mediated retrograde transport of TrkB (Fig. 2F-G and Suppl. Fig. 3C-D). Taken together, these data indicate that DCX decreases dynein-mediated retrograde transport through direct interactions with the dynein motor complex through its N-terminus, while the C-terminal domain of DCX negatively impacts this interaction to influence dynein-based cargo trafficking.

### The effects of DCX on retrograde trafficking require DCX-MT interactions

Previous work has shown that the binding of DCX to MTs contributes to DCX’s cellular functions (Moslehi et al., 2019; Reiner, 2013; Schaar et al., 2004; Yap et al., 2012) and that DCX’s MT interactions occur cooperatively (Bechstedt and Brouhard, 2012). To test whether MT binding and the underlying cooperativity of DCX’s MT interactions play a role in regulating dynein-based vesicular transport, we tested whether two DCX mutants, DCX A71S and T203R, could rescue the increase in retrograde transport of TrkB vesicles in DCX knockout neurons. These mutations, located in the R1 and R2 region of DCX, respectively, cause lissencephaly in humans and have been shown to decrease the cooperative MT binding of DCX (Bechstedt and Brouhard, 2012). Importantly, these mutations have no effect on DCX’s ability to associate with dynein *in vitro* (Suppl. Fig. 2D). Both mutants were unable to rescue the phenotype of increased retrograde transport of TrkB (Fig. 3A and Suppl. Fig. 4), suggesting that the cooperative binding of DCX to MTs is required for the DCX-induced decrease in dynein-based retrograde transport. In addition, our MT-binding assay demonstrates that, while C-DCX itself does not bind MTs, C-DCX increases the interactions of DCX with MTs (Fig. 3B). This suggests that the interactions between DCX and MTs are enhanced by DCX’s C-terminal domain, which is consistent with recent findings that the tail of DCX (amino acid 303 to the C-terminal end) helps to maintain the associations between DCX molecules on the MT lattice (Rafiei et al., 2022).

**Figure 3.**
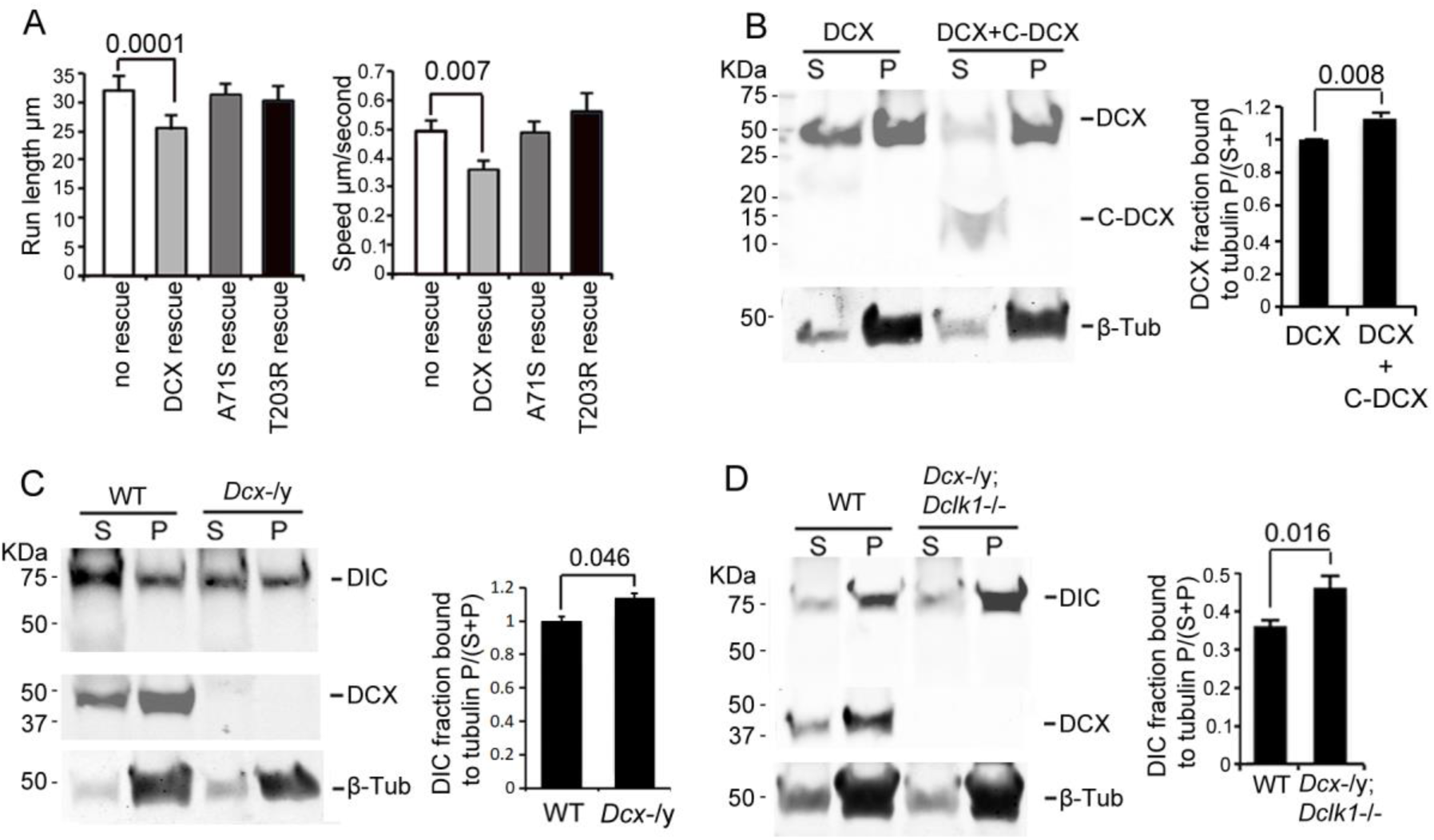
DCX association with MTs. (A). Dissociated cortical neuronal cultures from WT or Dcx-/y mice were transfected with plasmids expressing TrkB-RFP with/without plasmids expressing DCX-GFP, DCXA71S, or DCXT203R on DIV6 and imaged on DIV8. Quantification of Run length within 180 seconds and velocity are demonstrated. DCX, but not DCXA71S or DCXT203R, rescued the increased TrkB-RFP transport in DCX deficient axons. All quantifications are based on three independent experiments of each condition. P-values from t-tests are shown in each panel. Total numbers of neurons (N) and vesicles (V) used in the calculations are indicated in Suppl. Fig. 4. (B). Protein lysate from Hek293 cells expressing HA-DCX or HA-DCX plus HA-C-DCX are incubated with exogenously added MTs, which are then pelleted by ultracentrifugation. Western blot for HA in supernatant (S) or pellet (P) is performed to determine the amount of DCX or C-DCX associated with MTs. Representative Western blots are shown. Fraction of DCX bound to tubulin is calculated (Tubulin bound=P/(S+P)) and compared. Significantly more DCX is bound to MTs in the presence of C-DCX. P-value from t-test is shown. (C-D) Brain lysate from P0 WT, Dcx-/y or Dcx-/y;Dclk1-/- mice are incubated with exogenously added MTs, which are then pelleted by ultracentrifugation. Polymerized MTs are in the pellet. Western blot of DIC in supernatant (S) or pellet (P) is performed to determine the amount of DIC associated with MTs. Representative Western blots are shown. Fraction of DIC bound to tubulin is calculated (Tubulin bound=P/(S+P)) and compared between WT and Dcx-/y; or WT and Dcx-/y;Dclk1-/-. Significantly more DIC is bound to MTs in the absence of DCX. P-value from t-test is shown.

### DCX decreases dynein association with MTs

Since DCX enhances the binding of KIF1A to MTs and regulates KIF1A-mediated transport (Liu et al., 2012), we tested whether DCX also alters dynein’s interactions with MTs by performing a MT-binding assay using brain lysate from either WT, *Dcx-/y*, or *Dcx-/y;Dclk1-/-* mice. Our results show that significantly more DIC protein precipitates with MTs in the absence of DCX (Fig. 3C-D). Therefore, in contrast to DCX’s positive effects on KIF1A’s association with MTs (Liu et al., 2012), DCX decreases dynein-MT interactions.

### DCX negatively regulates dynein-mediated retrograde transport by regulating JIP3 association with dynein

Various binding partners are known to regulate dynein-based cargo transport (Vallee et al., 2012) and DCX could achieve its effect on the retrograde transport of TrkB by altering dynein’s association with regulatory proteins. Indeed, we previously reported that DCX regulates the interaction of dynein with its cargo adaptor JIP3 (Li et al., 2021), suggesting that DCX may regulate dynein-based cargo transport by controlling dynein- dynactin complex assembly and/or dynein-cargo attachment. In support of this idea, JIP3 also decreases the binding of DCX to dynein (Fig. 4A). As JIP3 serves as an adaptor protein for both kinesin and dynein (Arimoto et al., 2011; Drerup and Nechiporuk, 2013), DCX could differentially regulate anterograde vs. retrograde transport by competing with JIP3 for the binding of dynein. To test this possibility, we examined the retrograde transport of TrkB-RFP in *Dcx* knockout cortical neurons transfected with either control shRNA or JIP3 shRNA. Knockdown of JIP3 indeed significantly decreases the retrograde transport of TrkB (Fig. 4B-C, Suppl. Fig. 5 and Suppl. Video 5) while overexpression of JIP3 in WT neurons increases the retrograde transport TrkB (Fig. 4B-C, Suppl. Fig. 5 and Suppl. Video 6). Based on these results, we conclude that at least two mechanisms are at play when DCX regulates dynein-based transport: first, through negatively regulating dynein interactions with MTs, and secondly, through negatively regulating dynein interactions with JIP3.

**Figure 4.**
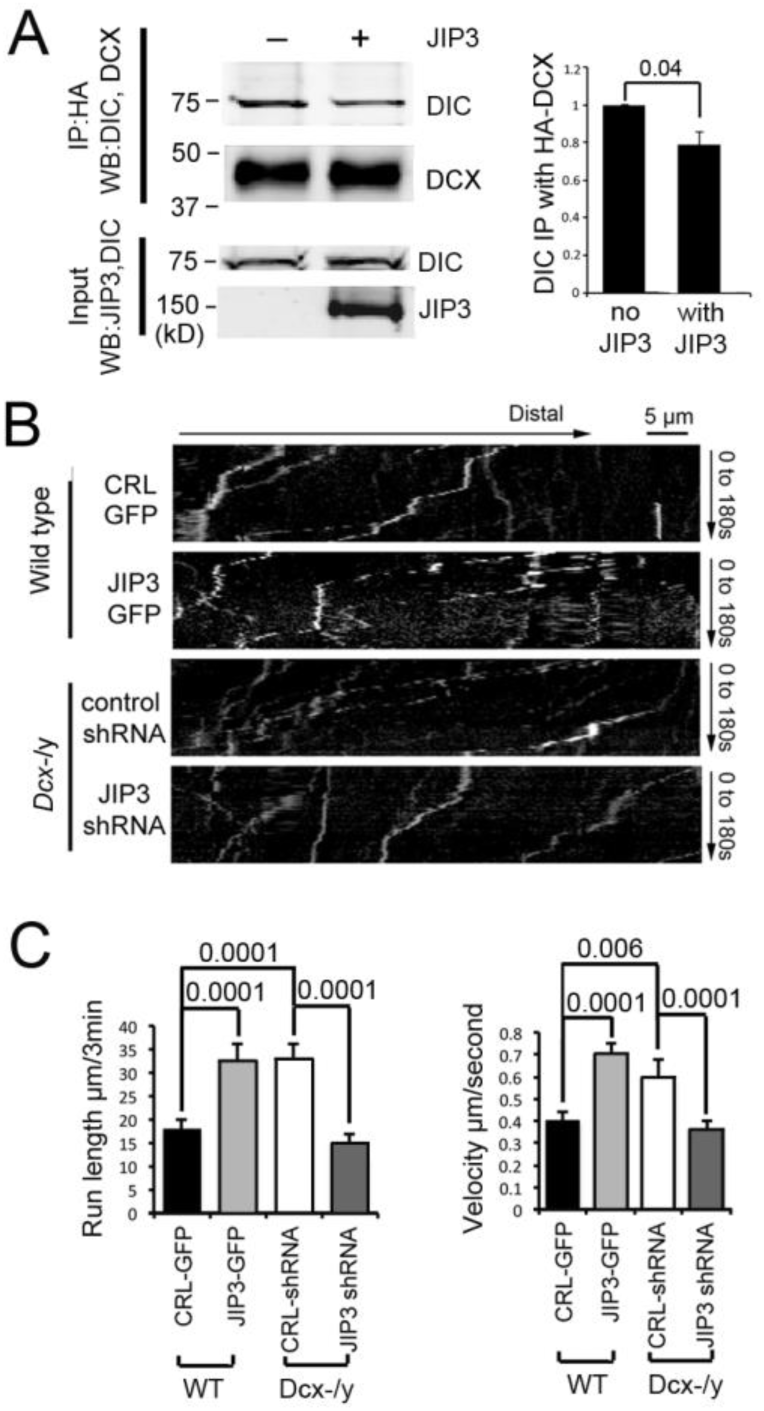
DCX and JIP3 competitively bind to the dynein motor complex, and JIP3 enhances retrograde transport mediated by dynein. (A) The resence of JIP3 decreases the interaction between DCX and DIC. HEK cells were transfected with plasmids expressing neuron-specific dynein intermediate chain isoform IC-1B and HA-tagged DCX with or without JIP3. Antibody for HA was used to precipitate HA-DCX and associated proteins. Western blot analysis of DIC and HA was performed to detect DIC immunoprecipitated with DCX. In the presence of JIP3, less DIC was associated with DCX, while total protein amount of either HA-DCX or DIC in the lysate were the same. Quantification of DIC bands of Western blot results (three independent experiments) after IP with HA were calculated and normalized with DIC levels in the lysate. P-value from t-test is shown. (B) Cultured cortical neurons from P0 WT mouse brains were transfected with plasmids expressing TrkB-RFP with or without JIP3-GFP. Neurons from Dcx-/y mouse brains were transfected with plasmids expressing TrkB-RFP with or without JIP3 shRNA. Representative kymographs of TrkB-RFP trafficking are demonstrated. (C) Quantification of TrkB run length and velocity. Overexpression of JIP3 significantly increases run length and velocity of TrkB in WT neurons. Downregulation of JIP3 by shRNA in Dcx-/y neurons decreases TrkB retrograde transport. P-values from t-tests are shown. All quantification data is based on three independent experiments of each condition. P-values from t-tests are shown in each panel. Total numbers of neurons (N) and vesicles (V) used in the calculations are indicated in Suppl. Fig. 5. See also Suppl. Video 5 and 6.

### Dynein, dynactin and JIP3 form a processive tripartite complex *in vitro*

To directly determine how DCX affects the motion of the dynein motor complex, we used total internal reflection fluorescence (TIRF) microscopy and performed single-molecule motility studies using purified components. Dynein, which assumes an auto- inhibited conformation in isolation (Torisawa et al., 2014; Zhang et al., 2017), moves processively along coverslip-attached MTs after its activation through the formation of a complex with its largest cofactor dynactin and a coiled-coil cargo adaptor protein such as Bicaudal D2 (BicD2) (McKenney et al., 2014; Splinter et al., 2012). Complexes such as dynein-dynactin-BicD2 (DDB), dynein-dynactin-BicDR1 (Bicaudal-D-related protein 1) (DDR), and dynein-dynactin-Hook3 (DDH), which have been recently shown to bind up to two dyneins (Grotjahn et al., 2018; Urnavicius et al., 2018), have been extensively studied using single-molecule TIRF assays (Christensen et al., 2021; McClintock et al., 2018; McKenney et al., 2014; Sladewski et al., 2018; Urnavicius et al., 2018). However, whether dynein-dynactin-JIP3 (DDJ) motor complexes can be reconstituted *in vitro* remains unknown.

Previous biochemical studies have shown that JIP3 interacts with dynein’s light intermediate chain (LIC) (Arimoto et al., 2011) and with kinesin-1’s light (Bowman et al., 2000) and heavy chains (Sun et al., 2011). Mutations in JIP3 result in the mis-localization of the dynein and impair retrograde transport (Arimoto et al., 2011; Celestino et al., 2021). However, the current consensus is that the coiled-coil region of JIP3 is too short to form a stable complex with dynein and dynactin (Chaaban and Carter, 2022; Lee et al., 2020; Reck-Peterson et al., 2018). Indeed, the putative N-terminal α-helix of JIP3, which is followed by an intrinsically disordered domain, extends only up to ∼180 residues according to the structural prediction by AlphaFold (Jumper et al., 2021) (Suppl. Fig. 6C). In contrast, cargo adaptors that have been successfully used for *in vitro* motility assays have been predicted to have substantially longer N-terminal coiled-coil domains. For example, the coiled coil of BicD2 is predicted to extend to ∼270 residues and cover the full shoulder of dynactin (Suppl. Fig. 6A-B), in agreement with cryo-EM studies (Urnavicius et al., 2015). Nonetheless, based on our *in vivo* and immunoprecipitation results, we hypothesized that JIP3 can form a tripartite complex with dynein and dynactin, possibly through a transition of its disordered domain into a more ordered conformation upon binding to dynein/dynactin. The latter idea is supported by the recently reported transition of a random coil in Nup358 into an α-helix upon binding to BicD2 (Gibson et al., 2022).

To test whether JIP3 can form an active DDJ motor complex, we generated and expressed a mouse JIP3 construct containing the N-terminal coiled-coil and the predicted adjacent intrinsically disordered domain (amino acids 1-240) in *E. coli* (Fig 5A and Suppl. Fig. 7A). The homolog of this construct in *Caenorhabditis elegans* has been shown to interact with dynein through the dynein LIC (Arimoto et al., 2011). To generate and purify full-length human dynein with its associated five subunits (IC, LIC, Tctex, LC8 and Robl), we co-expressed the five subunits with the HC of dynein in insect cells as done before (Suppl. Fig. 7C) (Schlager et al., 2014). To allow single-molecule fluorescence imaging of both dynein and JIP3, we labeled the dynein HC with SNAP-TMR via an N-terminal SNAP-tag and JIP3 with Halo-JF646 via a C-terminal HaloTag (Fig. 5A).

**Figure 5.**
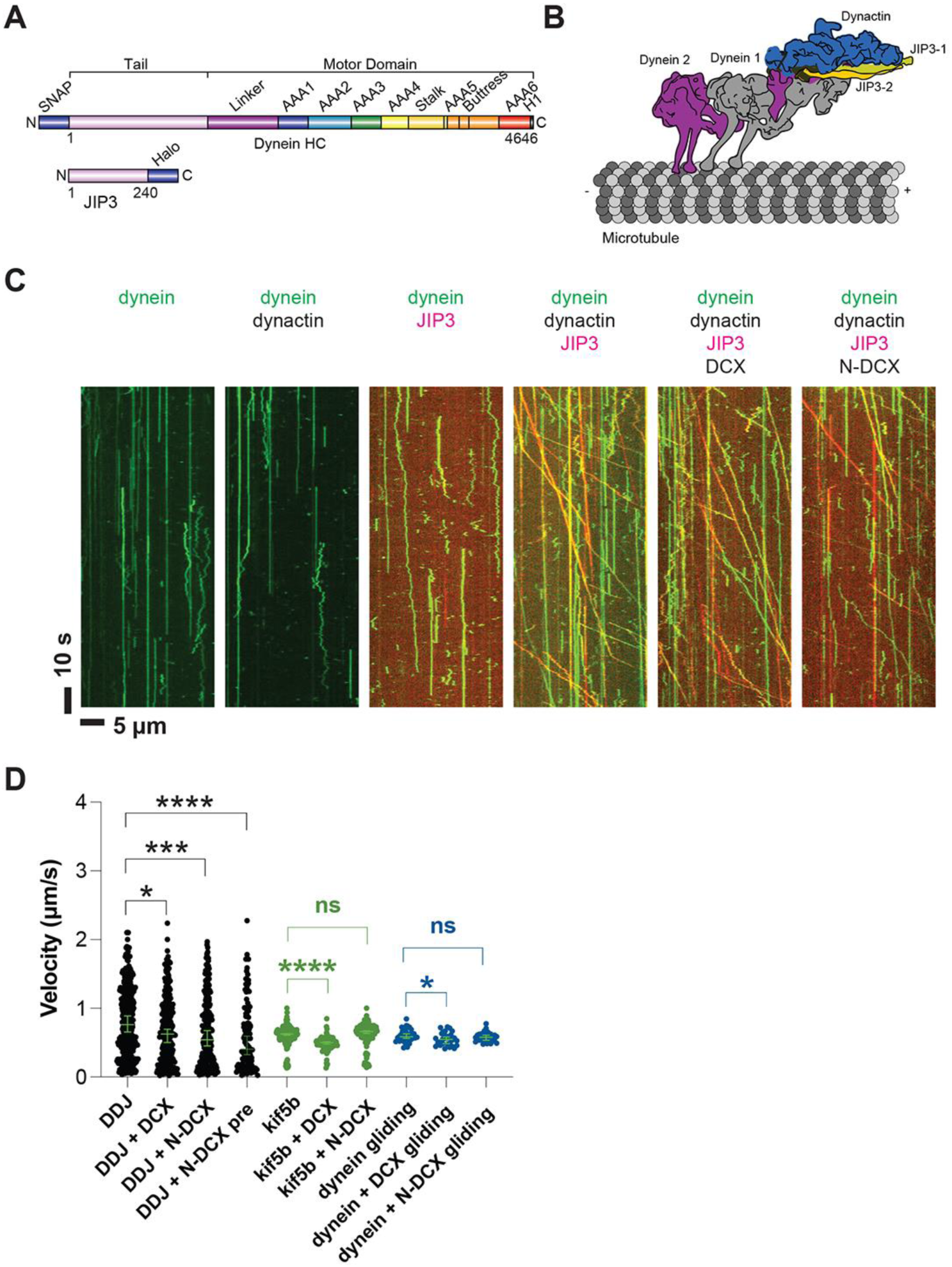
Dynein and dynactin form an active motor complex with JIP3 in vitro and DCX reduces its velocity. (A) Illustrations of the JIP3 and DCX constructs (left) and the DDJ motor complex (right). (B) Kymographs of dynein in the absence and presence of dynactin, JIP3, DCX and N-DCX. Dynein was labeled with SNAP-TMR (green) and JIP3 was labeled with Halo-JP646 (red). (C) The velocity of DDJ motor complexes, KIF5B, and gliding MTs powered by surface-absorbed single-headed dynein. The green bars represent the median with 95% CI. DDJ (DDJ only): 0.76 [0.65, 0.89] µm/s; DDJ + DCX (DDJ with 10 nM DCX): 0.62 [0.50, 0.70] µm/s (KS test, *p<0.1); DDJ + N-DCX (DDJ with 10 nM N-DCX): 0.54 [0.45, 0.67] µm/s (KS test, ***p<0.001); DDJ + N-DCX pre (dynein, dynactin, JIP3, and N-DCX assembled in the ratio of 1:1:1:1): 0.41 [0.32, 0.60] µm/s (KS test, ****p<0.0001). kif5b (kif5b only): 0.63 [0.61, 0.63] µm/s; kif5b + DCX (kif5b with 10 nM DCX): 0.50 [0.48, 0.51] µm/s (unpaired t -test, ****p<0.0001); kif5b + N-DCX (kif5b with 10 nM N-DCX): 0.66 [0.64, 0.67] µm/s (unpaired t -test, n.s.). MT gliding (powered by single-headed human dynein): 0.59 [0.56, 0.63] µm/s; dynein + DCX (MT gliding with 10 nM DCX): 0.55 [0.48, 0.58] µm/s (unpaired t -test, *p<0.1); dynein + N-DCX (MT gliding with 10 nM N-DCX): 0.58 [0.53, 0.61] µm/s (unpaired t -test, n.s.). From left to right, n = 342, 275, 252, 115, 234, 103, 117, 33, 31, 28. See also Suppl. Fig. 6-9, and Supple. Video 7.

As expected for an auto-inhibited motor, the purified dynein only diffused along MTs or bound rigidly (McKenney et al., 2014; Schlager et al., 2014), and the addition of dynactin (purified from cow brain (Schlager et al., 2014)), did not activate its motion (Fig 5B). While JIP3 transiently interacts with dynein (Suppl. Fig. 8A, Suppl. Video 7), it can neither stably bind to dynein (Suppl. Fig. 8B) nor activate dynein motion (Fig. 5B), similar to other studied dynein adaptors (McKenney et al., 2014; Schlager et al., 2014). However, strikingly, when we incubated JIP3 with dynein and dynactin on ice for 1 hour in a 1:1:1 stoichiometry, DDJ complexes formed and moved processively along MTs (Fig. 5B) at a velocity of 0.8 [0.7, 0.9] µm/s (median with 95% CIs) (Fig. 5C), which is comparable to other dynein complexes (Elshenawy et al., 2019; McKenney et al., 2014; Urnavicius et al., 2018). Our results demonstrate that JIP3 and dynactin are involved in dynein activation and that JIP3 can form highly processive DDJ complexes despite its predicted short coiled-coil domain.

### DCX decreases the velocity of DDJ motor complexes

To determine whether DCX negatively impacts the velocity of DDJ motor complexes as suggested by our *in vivo* results, we expressed full-length DCX and N-DCX with a C-terminal ybbR-tag (Yin et al., 2006) for labeling with CoA-CF488 or CoA-JF549 in *E. coli* (Suppl. Fig. 7B). We chose the small α-helical ybbR tag over the commonly used GFP tag (Bechstedt and Brouhard, 2012; Ettinger et al., 2016) to reduce possible steric blocking of dynein MT-binding sites by the introduced tag. At 10 nM concentration, DCX fully decorated MTs (Suppl. Fig. 9A) in the dynein motility buffer, while N-DCX had a much weaker affinity for MTs (Suppl. Fig. 9B), which is consistent with our immunoprecipitation data that demonstrated that the addition of C-DCX increased the affinity of DCX for MTs (Fig. 3B). Only when the ionic strength of the buffer was reduced, increasing amounts of N-DCX bound to MTs (Suppl. Fig. 9C). These observations confirm that our ybbR-tagged and labeled DCX constructs are functional.

Decorating MTs using 10 nM DCX slightly reduced DDJ’s velocity (0.62 [0.50, 0.70] µm/s, *p < 0.1) (Fig. 5C), while N-DCX had a stronger effect on the velocity (0.54 [0.45, 0.67] µm/s, ***p < 0.001) (Fig. 5C), which is consistent with our *in vivo* results (Fig. 2). The more pronounced effect of N-DCX on DDJ velocity supports our *in vivo* results that showed that the C-terminus of DCX negatively regulates DCX-dynein binding (Fig. 2E). Moreover, since N-DCX doesn’t bind MTs in regular motility buffer but only binds MTs in a motility buffer with half ionic strength (Suppl. Fig. 9B-C), N-DCX likely acts directly upon the DDJ complex rather than through MT binding.

To determine if N-DCX acts specifically on dynein, we also tested FL-DCX and N-DCX on another canonical MT-based motor, the kinesin-1 family member KIF5B. In contrast to the effects on DDJ, N-DCX has no effects on the velocity of KIF5B, while FL- DCX reduces the velocity of KIF5B (Fig. 5C), a result which agrees with a recent study that showed that DCX decreases the binding of kinesin-1 to MTs (Monroy et al., 2020). These results collectively suggest that DCX differentially affects the velocities of DDJ and kinesin-1: while DCX affects kinesin-1 motility through its binding to MTs, N-DCX affects DDJ motility through direct interactions with the dynein motor complex.

To dissect which component of the dynein motor complex N-DCX regulates, we first performed an MT-gliding assay using a N-terminal GFP-tagged single-headed human dynein construct expressed in insect cells (Htet et al., 2020). The single-headed dynein only contains the motor domain of the heavy chain, while the tail domain and all other subunits are absent. As this motor construct is non-processive (Trokter et al., 2012), we performed an MT-gliding assay to probe the effects of DCX on the activity of the dynein motor domain. To do so, we bound the GFP-tagged dyneins to a cover-glass surface via anti-GFP antibody at a motor-surface density that supports smooth gliding of MTs along the cover- glass surface. If N-DCX acts directly on the dynein motor domain as the dynein co-factor Lis1 does (Canty and Yildiz, 2020; DeSantis et al., 2017; Htet et al., 2020; Marzo et al., 2020), one would expect a reduction in gliding velocity when N-DCX is added. However, we found that N-DCX doesn’t affect MT gliding by single-headed dynein (Fig. 5C), which suggests that N-DCX regulates dynein through interactions with the dynein tail or through binding to dynein’s associated subunits. In contrast to N-DCX, we find that DCX decreases the MT gliding velocity slightly (Fig. 5C). This result implies that DCX also affects dynein-MT interactions through direct MT binding, while N-DCX, which does not bind MTs under the motility buffer conditions (Suppl. Fig. 9B), does not affect MT gliding powered by the dynein motor domain. This result implies that DCX may regulate dynein function via two pathways: through direct interactions with the dynein motor complex and through the binding to MTs.

### DCX interferes with the recruitment of a second dynein to DDJ

Previous studies have demonstrated that BicD2 can recruit two dimeric dyneins and that a DDB complex with two dyneins shows higher velocities compared to a DDB complex with one dynein (Elshenawy et al., 2019; Sladewski et al., 2018; Urnavicius et al., 2018). Adaptors such as BicDR1 and Hook3, which predominantly recruit two dyneins, also show increased velocities compared to DDR and DDH complexes with only one dynein (Elshenawy et al., 2019; Urnavicius et al., 2018). We note that the velocity reduction of DDJ in the presence of N-DCX is similar to the velocity reduction when a two-dynein motor assembly loses a dynein motor. Moreover, when we assembled DDJ complexes in the presence of N-DCX, the velocity of DDJ was reduced further (0.41 [0.32, 0.60] µm/s, ****p<0.0001) (Fig. 5C).

To test the hypothesis that N-DCX displaces the second dynein from a DDJ motor complex with two dyneins, we first determined whether JIP3 permits the recruitment of two dyneins. To do so, we assembled DDJ complexes using equal amounts of dynein labeled with TMR and Alexa-647. In the absence of DCX, we indeed observed moving DDJ complexes with two colocalized colors, demonstrating that DDJ can recruit two dyneins (Fig. 6A). The fraction of colocalization was 42 ± 2% (mean ± SEM), which is comparable to the colocalization of DDR and DDH complexes with two differently labeled dyneins (Elshenawy et al., 2019; Urnavicius et al., 2018). In support of our hypothesis that N-DCX displaces a dynein from a two-dynein motor complex, addition of N-DCX reduced the co-localization fraction to 27 ± 3% (Fig. 6B), which is close to the reported colocalization of the DDB motor complex (Elshenawy et al., 2019). In addition, those few remaining DDJ complexes that contained two colocalized dyneins despite the presence of N-DCX, showed a reduced velocity in the presence of N-DCX (Fig. 6C). In conclusion, similar to DDR and DDH, DDJ predominantly recruits two dyneins, and N-DCX interferes with the binding of the second dynein; N-DCX can still affect the velocity of the DDJ complex with even two dyneins, possibly via disrupting interaction between the tails of the two dyneins (Elshenawy et al., 2019).

**Figure 6.**
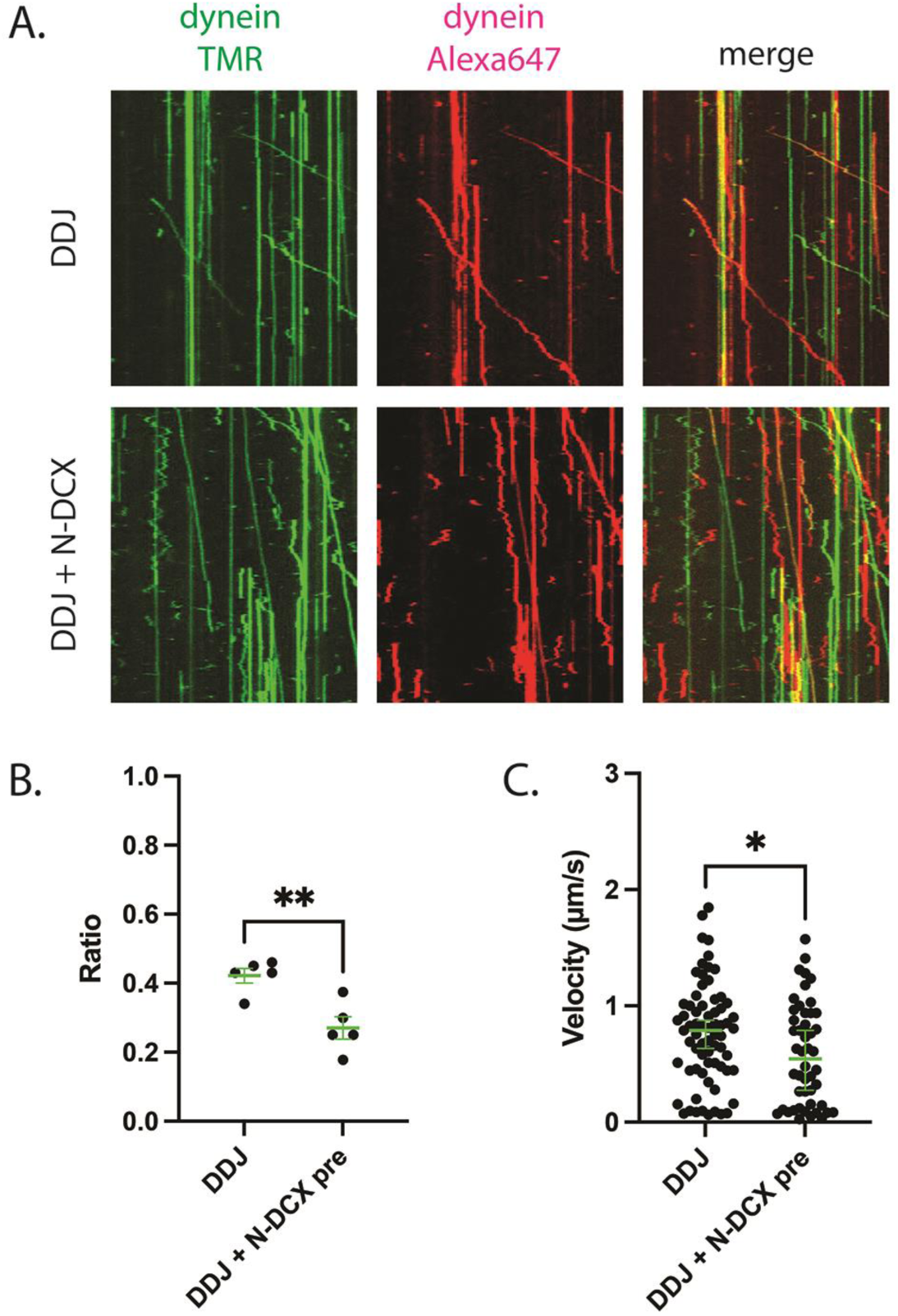
DDJ motor complexes associate with two dyneins and N-DCX negatively affects its velocity by displacing the second dynein. (A) Kymograph of DDJ assembled in the absence (top) or presence (bottom) of N-DCX with dynein that were labeled separately with SNAP-TMR and SNAP-Alexa647. (B) The ratio of two-color moving molecules versus the total moving molecules. The green bars represent mean ± SEM. DDJ: 42 ± 2%; DDJ + N -DCX pre: 27 ± 3% (unpaired t -test, **p<0.01). The molecules within each field of view were counted to produce a single value (50 µm x 50 µm). (C) The velocity of two -color moving molecules. The green bars represent median with 95% CI. DDJ: 0.79 [0.63, 0.87] µm/s; DDJ + N- DCX pre: 0.54 [0.27, 0.79] µm/s (KS-test, *p<0.1).

### Rescuing retrograde transport defects in *Dcx-/y; Dclk1-/-* neurons ameliorates neuronal migration defects

One of the characteristics of DCX-linked lissencephaly is a profound defect in cortical neuronal migration. We therefore asked whether the effects of DCX on dynein-based retrograde transport we observe play a role in the migration of cortical neurons during development. If the answer is yes, rescuing the abnormally increased dynein-based retrograde trafficking should mitigate the cortical neuronal migration defects observed in the developing *Dcx-/y; Dclk1-/-* mouse brain (Deuel et al., 2006; Koizumi et al., 2006). Since cortical neuronal migration is relatively normal in the *Dcx-/y* mouse (Corbo et al., 2002), we used a *Dcx-/y; Dclk1-/-* mouse, which has a cortical neuronal migration defect as the Dcx-redundant gene Dclk1is knocked out as well (Deuel et al., 2006; Koizumi et al., 2006). A plasmid expressing GFP and a shRNA that specifically targets DHC (Tsai et al., 2007) was micro-injected into the lateral ventricle of embryonic day (E)14.5 *Dcx-/y;Dclk1-/-* mouse brains and transfected using in utero electroporation. Mouse embryos were then sacrificed on E18.5. As expected, down-regulating DHC partially rescued the retention of neuroblasts in the deeper region of the cortex observed in *Dcx-/y; Dclk1-/-* mouse brains (Fig. 7A-B). Based on these results, we wondered whether the dysregulation of dynein is in part due to increased association of JIP3 with dynein in the absence of DCX and whether downregulation of JIP3 expression may also ameliorate neuronal migration defects. To test this possibility, we microinjected plasmids expressing JIP3 shRNA1 and GFP into the lateral ventricle of E14.5 *Dcx-/y; Dclk1-/-* embryos and transfected the plasmids into neural progenitors using in utero electroporation. In agreement with our hypothesis, down regulation of JIP3 in *Dcx-/y; Dclk1-/-* mouse brain significantly rescued the lamination defect (Fig. 7C-D). Collectively, our results demonstrate the importance of the regulation of dynein-dependent retrograde trafficking by DCX and JIP3 during neuronal migration.

**Figure 7.**
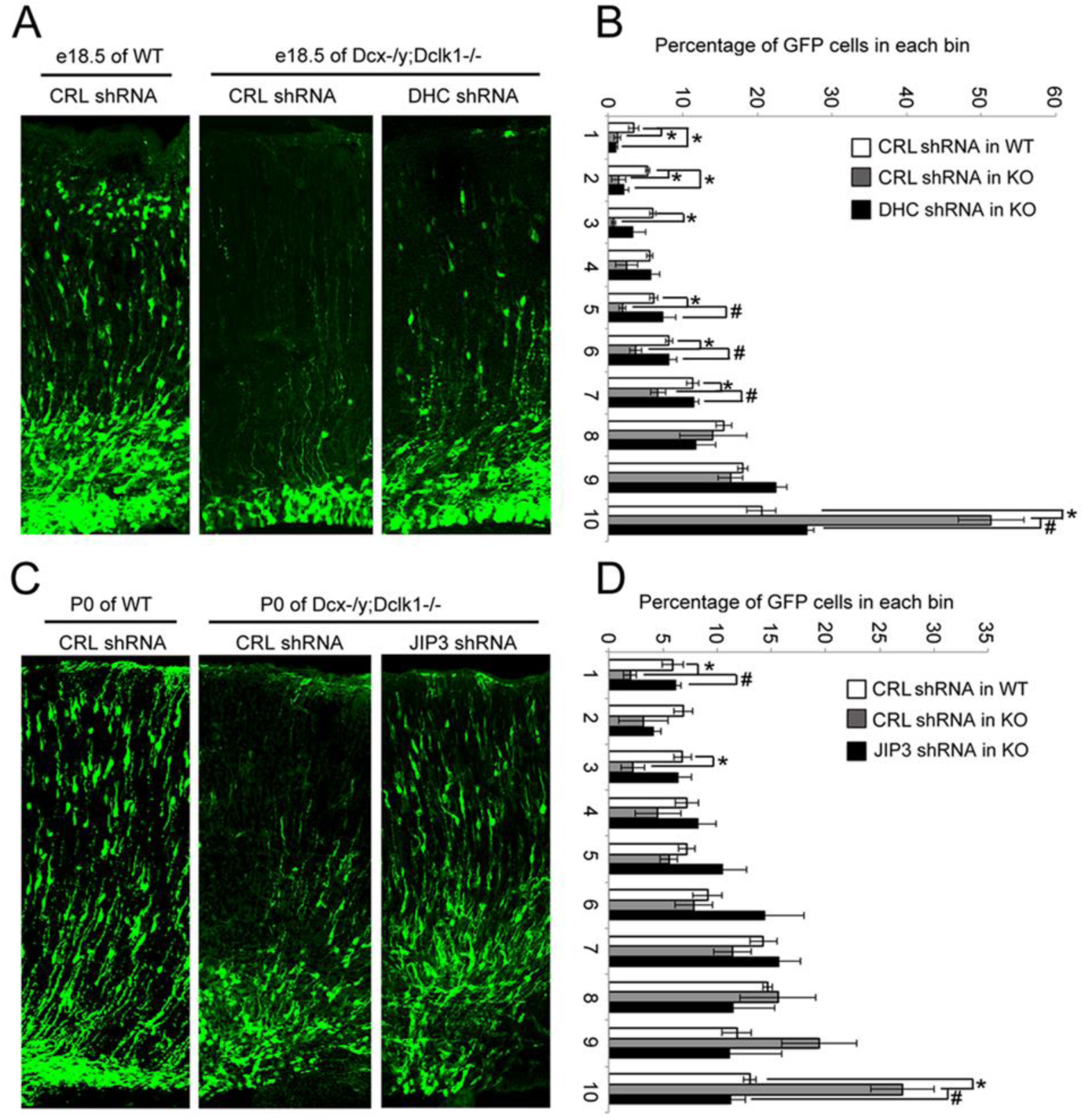
Knock-down of DHC or JIP3 in *Dcx-/y; Dclk1-/-* in mouse cortex partially rescues the defect of pyramidal cell migration. (A) GFP-positive neurons were imaged and counted at E18.5 after electroporation at E14.5 with vectors expressing control (CRL) shRNA (+GFP) or DHC shRNA (+GFP). (B) Percent of GFP positive cells in evenly divided regions of the cortex (1-10) from the pia to the lateral ventricle. Asterisks denote statistically significant p-values (t-test, p< 0.05) between WT with CRL shRNA and *Dcx-/y; Dclk1-/-* with CRL shRNA. Number sign (#) denotes p<0.05 of t-test between *Dcx-/y; Dclk1-/-* with CRL shRNA and *Dcx-/y; Dclk1-/-* with DHC shRNA. The data represent the mean±SEM of three different brains in each condition. (C) GFP-positive neurons were imaged and counted at P0 after electroporation at E14.5 with vectors expressing control (CRL) shRNA (+GFP) or JIP3 shRNA (+GFP). (D) Percent of GFP positive cells in evenly divided regions of the cortex (1-10) from the pia to the lateral ventricle of different mouse brains. Asterix (*) denotes p<0.05 of t-test between WT with CRL shRNA and *Dcx-/y; Dclk1-/-* with CRL shRNA. Number sign (#) denotes p<0.05 of t-test between *Dcx-/y; Dclk1-/-* with CRL shRNA and *Dcx-/y; Dclk1-/-* with JIP3 shRNA. The data represent the mean±SEM of three individual brains in each condition.

## DISCUSSION

Previous reports have linked DCX, a causative gene for classical lissencephaly in males, to defects in dynein-based functions in neurons (Kaplan and Reiner, 2011; Li et al., 2021; Tanaka et al., 2004), but how DCX modulates dynein functions has remained unclear. In this study, we demonstrate for the first time that DCX negatively regulates dynein-mediated retrograde trafficking in neuronal axons through its interactions with MTs and through interactions with the dynein motor complex (Fig. 8). We show that DCX decreases the velocity and processivity of dynein-based cargo transport *in vivo* and the velocity of dynein-dynactin-JIP3 motor complexes *in vitro* and demonstrate that the DCX-based regulation of dynein-driven retrograde transport is important to cortical development. Combined with our previous finding that DCX positively regulates KIF1A-mediated anterograde transport (Liu et al., 2012), we conclude that DCX differentially regulates anterograde and retrograde intracellular trafficking in neuronal axons, and therefore mediates the transport of critical protein complexes during neuronal growth and development.

**Figure 8.**
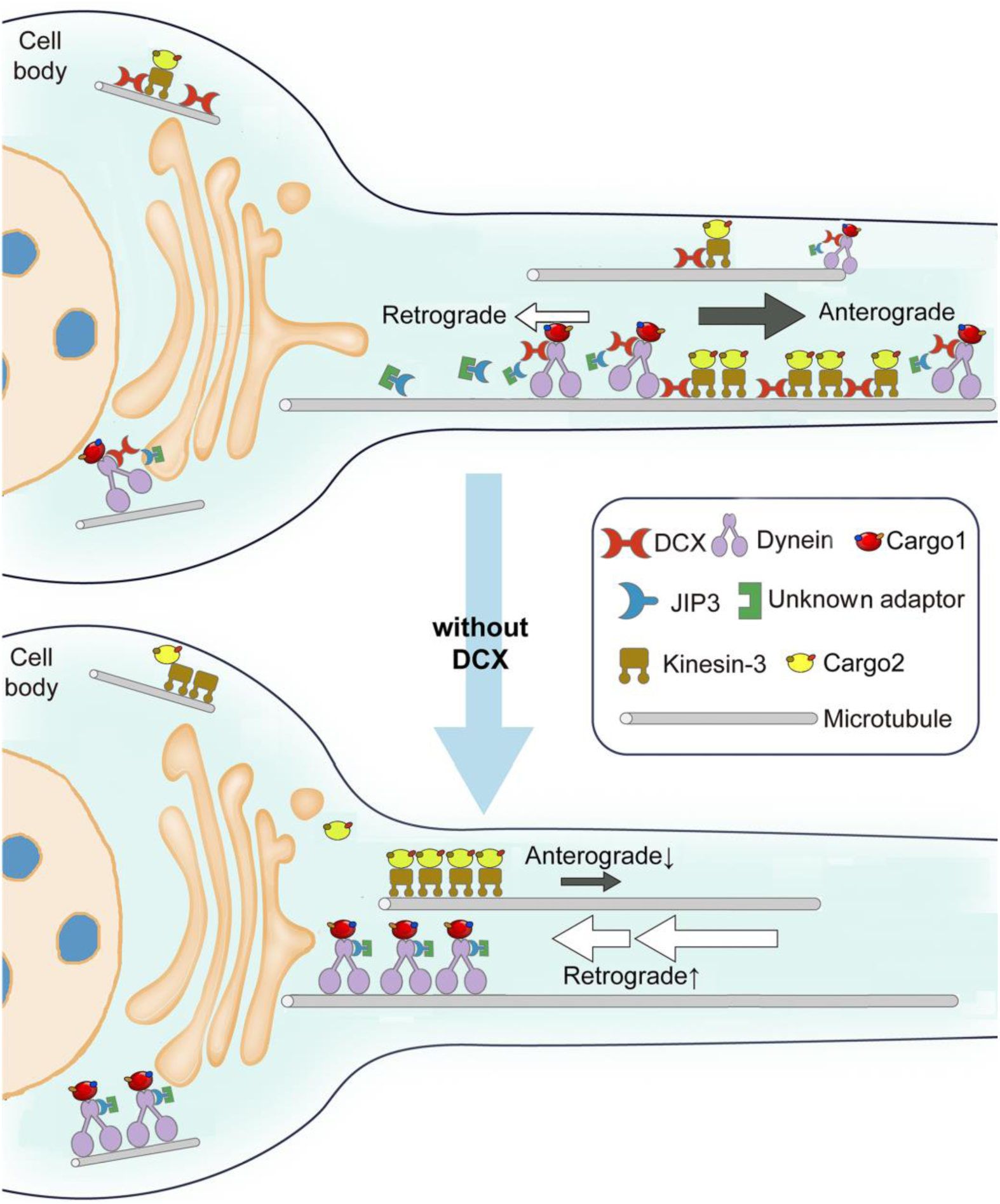
Schematic diagram shows the regulation of dynein-mediated retrograde transport by DCX. Cargo-bound dynein motor complex drives retrograde transport from plus end of MTs (distal axon) to minus end of MTs (cell body). In WT neurons, DCX association with kinesin-3 helps kinesin–3- mediated anterograde transports (Liu et al., 2012). DCX decreases dynein-MT interactions (represented by tilted dynein complex along MT). DCX and JIP3 competitively associate with dynein. When DCX binds dynein, very few JIP3 proteins associate with dynein, the retrograde transport is normal. In DCX KO neurons (without DCX), Kinesin–3-mediated anterograde transports are decreased without DCX (Liu et al., 2012). Meanwhile, more JIP3 molecules bind dynein, which also associates with MT stronger without DCX. The dynein mediated retrograde transport is faster. The balance between anterograde transport and retrograde transport is broken without DCX.

### DCX negatively regulates dynein motion through interactions with both dynein and MTs

Studies from other labs demonstrated that kinesin and dynein engage in a “tug-of-war” when attached to the same cargo (Belyy et al., 2016; Gennerich and Schild, 2006; Rezaul et al., 2016). It is therefore possible that DCX’s effects on dynein-mediated retrograde transport are indirect effects through anterograde transport. However, our data show that DCX affects retrograde transport directly, both through its binding to MTs and through its direct interactions with the dynein motor complex. The importance of DCX-MT interactions for retrograde transport is based on our observation that the pathogenic DCX mutations A71S and T203R, which decrease the cooperative binding of DCX to MTs but have no effect on DCX and dynein interactions, fail to restore dynein-mediated retrograde trafficking. At the same time, our data reveal that DCX directly interacts with the dynein motor complex through its N-terminal domain both *in vivo* and *in vitro*, and this interaction also negatively regulates retrograde transport. Our immunoprecipitation experiments show that N-DCX without DCX’s C-terminal domain has a stronger affinity for dynein, while our *in vivo* results indicate that N-DCX has a stronger inhibitory effect on dynein-mediated retrograde transport than the FL-DCX; these results are further supported by our *in vitro* experiments that demonstrate that N-DCX more strongly reduces the velocity of reconstituted DDJ complexes than FL-DCX. Interestingly, DCLK1 interacts with KIF1A also through its N-terminal domain (Lipka et al., 2016). In summary, interactions between DCX and dynein influence dynein-mediated retrograde transport.

Our data further demonstrate that DCX’s C-terminal S/P-rich domain decreases DCX-dynein association, although C-DCX itself does not interact with either dynein or MTs. Since the DCX C-terminus has several phosphorylation sites (Graham et al., 2004; Jin et al., 2010; Shmueli et al., 2006; Slepak et al., 2012; Tanaka et al., 2004), it will be interesting in future studies to determine whether phosphorylation of residues in DCX’s C-terminus regulates the association of DCX with dynein.

### DCX regulates dynein-mediated retrograde transport through JIP3

DCX association with dynein also alters the composition of the dynein motor complex. In our previous study, we found that the presence/absence of DCX most strongly altered the amount of the signaling adaptor protein, JIP3, that immunoprecipitated with the dynein motor complex (Li et al., 2021). In this study, we find that DCX and JIP3 competitively associate with dynein and that a DCX-induced reduction in the association of JIP3 with dynein results in diminished dynein-mediated retrograde transport. Thus, when DCX is absent, more dynein motors associate with MTs, and more JIP3 associates with dynein—events that greatly promote retrograde trafficking.

Our previous work also showed that the effects of DCX on the dendritic localization and patterning of the somatic Golgi apparatus depend on JIP3 and dynein (Li et al., 2021). Since the relocation of the Golgi apparatus from the soma to dendrites occurs along MTs, DCX could promote this process by upregulating anterograde and downregulating retrograde trafficking through its activating effects on KIF1A and its inhibiting effects on dynein/JIP3, respectively.

JIP3 belongs to the JIP family of proteins, which interact with C-Jun N-terminal Kinase (JNK). All mammalian JIP proteins are expressed in the brain (Dickens et al., 1997; Ito et al., 1999; Kelkar et al., 2000; Kelkar et al., 2005; Yasuda et al., 1999). Sunday Driver, the JIP3 homolog in *Drosophila*, directly binds to kinesin-1 (Bowman et al., 2000; Byrd et al., 2001; Sun et al., 2011). UNC16, the JIP3 homolog in *C. elegans*, interacts with both kinesin-1 and dynein (Byrd et al., 2001). JIP3 colocalizes with the dynein-dynactin motor complex and serves as an adaptor protein for dynein-mediated retrograde transport of active JNK and lysosomes (Cavalli et al., 2005; Drerup and Nechiporuk, 2013).

The targeted deletion of JIP3 has a similar phenotype to that of *DCX -/y; Dclk1-/-* mice (Deuel et al., 2006; Fu et al., 2013; Koizumi et al., 2006) with disrupted formation of the telencephalon and the agenesis of the telencephalic commissures, possibly through impaired vesicle transport and defects in axon guidance (Ha et al., 2005; Kelkar et al., 2003). Furthermore, previous studies showed that JIP3 regulates axon branching through GSK3β-signaling pathway by phosphorylation of DCX at Ser327, which is located at C-terminal S/P-rich region of DCX (Bilimoria et al., 2010). Therefore, JIP3 might regulate the association of DCX with dynein through phosphorylation. Since JIP3 is also involved in kinesin-based transport, it may be a candidate for mediating cross-talk between anterograde and retrograde motors. A previous study indicates that DCX interacts with another JIP family protein, c-Jun N-terminal kinase (JNK) interacting protein (JIP1). The phosphorylation of DCX by the JNK pathway is important for neuronal migration (Gdalyahu et al., 2004). It will be interesting to explore in future studies whether the JIP1/JNK pathway is involved in DCX effects on dynein functions.

### DCX regulates the assembly and motility of the dynein-dynactin-JIP3 motor complex

By reconstituting for the first time the *in vitro* motility of DDJ motor complexes and by demonstrating—using two-color single-molecule co-localization studies—that DDJ associates with up to two dyneins, we have revealed that DCX competes with the binding of the second dynein to DDJ, resulting in reduced velocities of the moving motor complexes. While numerous dynein-dynactin-adaptor complexes such as DDB, DDR and DDH have been extensively studied *in vitro* using single-molecule TIRF microscopy (Christensen et al., 2021; McClintock et al., 2018; McKenney et al., 2014; Sladewski et al., 2018; Urnavicius et al., 2018), it was the consensus that the predicted ∼180 amino acids α- helical coiled-coil region in JIP3 is too short to be capable of forming a tripartite complex with dynein and dynactin (Chaaban and Carter, 2022; Lee et al., 2020; Reck-Peterson et al., 2018). Surprisingly, however, we found that a truncated JIP3 construct containing the N-terminal coiled-coil and the predicted adjacent intrinsically disordered domain (amino acids 1-240) can form an active DDJ complex with two dyneins. N-DCX reduced the velocity of DDJ complexes with two dyneins from ∼0.8 µm/s to ∼ 0.4 µm/s as a result of the dissociation of the second dynein. That N-DCX’s inhibitory effect on DDJ is via interacting with dynein’s tail domain or dynein’s associated subunits, but not with dynein’s motor domain or the MTs. The conclusion is supported by the findings that N-DCX, which does not bind MTs under our assay condition, does not impact the MT-gliding activity by a tail-truncated single-head dynein (a recombinant construct that contains the motor domain and the linker), while FL-DCX, which decorates MTs well, reduces the MT-gliding velocity slightly.

The fact that DDJ complexes are active and associate with two dyneins, implies that either the predicted disordered region in JIP3 following the first α-helix forms an α-helical structure (possibly when JIP3 interacts with dynactin and the dynein tail) or that DDJ associates with two JIP3 molecules as has been recently shown for BICDR1 and Hook (Chaaban and Carter, 2022). This result contrasts with a previous report that demonstrated that while a short Hook3 construct could form a stable complex with dynein and dynactin, the resulting DDH complex was incapable of moving along MTs (Schroeder and Vale, 2016). It is possible that Hook3 and JIP3 differently interact with dynactin and dynein, which could result in a different degree of activation. Indeed, cryo-EM studies show that different adaptors bind to dynactin and dynein differently (Urnavicius et al., 2018). These findings collectively suggest that cargo adaptors fine-tune dynein’s activity by utilizing different interactions with dynein and dynactin. While we also found that a longer JIP3 construct (aa 1-548) formed an active complex with dynein and dynactin (data not shown), this construct was prone to aggregation. Of note, full-length BicD2 has been shown to be autoinhibited by its third coiled-coil domain (Hoogenraad et al., 2001; McClintock et al., 2018). It is therefore possible that also our longer JIP3 construct is more likely to be autoinhibited. We therefore used the shorter JIP3 construct for our studies. Collectively, our *in vitro* reconstitution studies with purified proteins agree with our *in vivo* observations that DCX downregulates dynein’s activity, and that the C-terminus of DCX auto-inhibits DCX’s interaction with dynein. Indeed, a recent study has shown that the C-terminus of DCX facilitates the binding of neighboring DCX molecules to MTs via intermolecular interactions with DCX’s N-terminal domain (Rafiei et al., 2022), which suggests that the C- and N-terminal domains of DCX have an intrinsic affinity for each other.

### How does disinhibition of dynein by DCX-based loss-of-function lead to defects in early neuronal development?

Loss of function mutations in dynein and its co-factors can cause malformations of cortical development (Feng and Walsh, 2004; Pawlisz et al., 2008; Poirier et al., 2013; Reiner et al., 1993; Sasaki et al., 2005; Youn et al., 2009). Our study shows for the first time that abnormally increased dynein function, as observed in mice with DCX knockout, can also cause defects in development resulting in cortical malformations. Our neuronal migration studies performed by *in utero* electroporation, shows that diminishing dynein activity by either knocking down DHC or by decreasing the amount of JIP3 can partially rescue the defects in *Dcx-/y;Dclk1 -/-* mouse cortex. The fact that the rescue is regional and in the deepest regions of the cortex near the ventricular and subventricular zone implies that early defects in neural progenitor or neuroblast biology may be preferentially affected by increased dynein activity in mice with lacking DCX. When dynein activity is abnormally and globally increased, precise spatial regulation of motor function is lost in neural progenitors and neurons, which may have effects on cell-biological functions mediated by dynein, including progenitor cell division, nucleokinesis, and polarized transport of signaling molecules (Roberts et al., 2013; Tsai et al., 2010; Vale, 2003).

Therefore, loss of dynein inhibition may have direct effects on important signal transduction pathways from distal neuronal processes, including neurotrophin (BDNF) (Bhattacharyya et al., 2002) and mitogen activated protein kinase signaling via JNK (Rishal and Fainzilber, 2014). JIP3 is known to bind dynein upon JNK activation and is, thus, an important mediator of the mitogen activated protein kinases (Drerup and Nechiporuk, 2013). While JIP3’s effects on BDNF signaling have not been appreciated previously, JIP3 enhanced retrograde transport of the canonical BDNF cargo, TrkB, is consistent with the known cross-talk between neurotrophic and mitogen-activated protein kinase signaling (Huang et al., 2011). Alternatively, JIP3 may be a more general dynein cofactor for mediating retrograde trafficking. While DCX’s effects on dynein and JIP3 occur predominantly in development, the activity of other DCX-family proteins, which are expressed in mature neurons (Reiner et al., 2006), have important consequences for understanding neuron-specific signaling that extend beyond development to degeneration, injury, and repair.

Dynein and KIF1A regulate apically-directed or basally-directed nuclear movement, respectively, of radial glial progenitor cells (Tsai et al., 2010). It is suggested that DCX mediates KIF1A’s effect on basally-directed nuclear movement through the BDNF pathway (Carabalona et al., 2016), while influencing dynein’s role in apically-directed nuclear movement through regulating the perinuclear MT structure (Tanaka et al., 2004). Based on our results in this study, DCX might also regulate nuclear migration through influencing the balance of KIF1A/dynein-mediated anterograde/retrograde transport via its regulation of JIP3 association with the two motors. Further studies are needed to prove this.

## EXPERIMENTAL PROCEDURES

### Antibodies and Reagents

Cell culture reagents were purchased from Life Technologies (Grand Island, NY). Antibodies to DCX (ab18723), DHC (ab6305), DIC (ab23905), and Tubulin (ab6161) are purchased from Abcam (Cambridge, MA). Antibody to JIP3 is from Santa Cruz Biotechnology (Dallas, TX). Antibody to HA is from EMD Millipore (Billerica, MA).

Construct expressing IC-1B is generously provided from Dr. Kevin Pfister (UVA). Construct expression JIP3 is generously provided from Dr. Roger Davis (UMASS MED). Construct expression TrkB-RFP is generously provided from Dr. Xiaowei Zhuang (Harvard University). All other reagents are purchased from Sigma-Aldrich (St. Louis, MO).

### Mammalian expression and RNA interference constructs

DNA sequences for HA-tagged N-DCX (1-270 N-terminal amino acids) and C- DCX (271-361 amino acids) are synthesized by PCR using construct expressing FL-DCX (Liu et al., 2012) as template, and then cloned into plasmid pBA (Jacobson et al., 2006). HA-tagged DCX mutant T203R were created using QuikChange Site-Directed Mutagenesis kit (Stratagene). HA-tagged DCX mutant A71S was synthesized commercially (Genewiz) and subcloned into plasmid pBA. All RNAi control or target sequences (hp) were cloned into the pSilencer 1.0-U6 plasmid. The complementary RNAi oligos were annealed and ligated into pSilencer-GFP (gift from Shirin Bonni) (Sarker et al., 2005).

### Animals and Primary Cortical Neuron Cultures

All animal procedures were approved by the Committee on the Ethics of Animal Experiments of Wenzhou Medical University (#wydw2019-0723). P0 cortices were dissected and dissociated using the Worthington papain dissociation system (Worthington Biochemical Corp., Lakewood, NJ). Neurons were plated on Poly-L-ornithine solution coated coverglasses in neuronal culture medium (Neurobasal medium plus B27, Glutamine, FGF (10ug/ml) and Pen/Strep) until experiments.

### Time-Lapse Imaging

Cultured cortical neurons were transfected with different constructs on DIV6 using Lipofectamine 2000 according to manufacturer’s instruction. Images were acquired on an inverted epifluorescence microscope (IX-81, Olympus America Inc., Melville, NY) equipped with high numerical aperture lenses (Apo 603 NA 1.45, Olympus) and a stage top incubator (Tokaihit, Japan) maintained at 37 °C at a rate of one capture per 3 seconds. Fluorescence excitation was carried out using solid-state lasers (Melles Griot, Carlsbad, CA) emitting at 488 nm (for green) and 561 nm (for red) fluorophores. Emission was collected through appropriate emission band-pass filters obtained from Chroma Technologies Corp. (Brattleboro, VT). Images were acquired with a 12-bit cooled CCD ORCA-ER (Hamamatsu Photonics) with a resolution of 1280 3 1024 pixels (pixel size = 6.45 mm^2^). The camera, lasers, and shutters were all controlled using Slidebook 5 (Intelligent Imaging Innovations, Denver, CO). For all calculations and measurements of vesicle movement, only bright vesicles located in the proximal region of axons (∼100 µm away from cell body) are analyzed. A vesicle is counted as mobile only if the displacement is at least 5 µm. A vesicle is counted as stationary if moves less than 5 µm. To calculate the run length and velocity, vesicles were analyzed only if the net run length is at least 5 µm in retrograde direction. The velocity is calculated as: the length of a continuous retrograde movement divided by the length of the time. Those stationary vesicles are not counted for velocity. Analysis of timelapse imaging was performed with MetaMorph for tracking and the ImageJ Manual Tracking plugin as described (http://rsbweb.nih.gov/ij/plugins/track/track.html).

### Pull-down Assay and Mass Spectrometry procedure and analysis

HA or HA-tagged DCX proteins were immobilized on Anti-HA agarose beads and subsequently mixed with protein lysates from embryonic day-18 mouse brains and incubated with rotation for 16 h at 4 °C to pull down associating proteins. The beads were washed four times. The beads were then incubated with DTT solution (final concentration of 10 mmol/L) and reduced in a 56 °C water bath for 1 h. IAA solution was added (final concentration of 50 mmol/L) and protected from light for 40 min. The proteins were digested with trypsin overnight at 37°C. After digestion, the peptides were desalted using a desalting column, and the solvent was evaporated in a vacuum centrifuge at 45 °C. The peptides were dissolved in sample solution (0.1% formic acid in water) and ready for mass spectrometry analysis. Samples were loaded onto Nanocolumn (100 μm×10 cm) packed with a reversed-phase ReproSil-Pur C18-AQ resin (3 μm, 120 Å, Dr. Maisch GmbH, Germany). The mobile phases consisted of A (0.1% formic acid in water) and B (acetonitrile). Total flow rate is 600 nL/min using a nanoflow liquid chromatograph (Easy- nLC1000, ThermoFisher Scientific, USA). LC linear gradient: from 4% to 8% B for 2 min, from 8% to 28 % B for 43 min, from 28 % to 40% B for 10 min, from 40% to 95% B for 1 min and from 95% to 95% B for 10 min. Eluted peptides were introduced into the mass spectrometer (Q Exactive™ Hybrid Quadrupole-Orbitrap™ Mass Spectrometer, Thermo Fisher Scientific, USA). The spray voltage was set at 2.2 kV and the heated capillary at 270°C. The machine was operated with MS resolution at 70000 (400 m/z survey scan), MS precursor m/z range: 300.0-1800.0. The raw MS files were analyzed and searched against protein database based on the species of the samples using MaxQuant (1.6.2.10). The parameters were set as follows: the protein modifications were carbamidomethylation (C) (fixed), oxidation (M) (variable), Acetyl (Protein N-term) (variable); the enzyme specificity was set to trypsin; the maximum missed cleavages were set to 2; the precursor ion mass tolerance was set to 20 ppm, and MS/MS tolerance was 20 ppm. Only high confident identified peptides were chosen for downstream protein identification analysis. RIPA Lysis and Extraction Buffer, Pierce™ BCA Protein Assay Kit were purchased from Thermo Fisher Science. DL-dithiothreitol (DTT), iodoacetamide (IAA), formic acid (FA), acetonitrile (ACN), were purchased from Sigma (St. Louis, MO, USA), trypsin from bovine pancreas was purchased from Promega (Madison, WI, USA). Ultrapure water was prepared from a Millipore purification system (Billerica, MA, USA). An Ultimate 3000 system coupled with a Q Exactive™ Hybrid Quadrupole-Orbitrap™ Mass Spectrometer (Thermo Fisher Scientific, USA) with an ESI nanospray source.

### Microtubule-binding assay

Mouse brains are dissected and flash frozen and kept at -80 °C until experiment. Flash frozen mouse brains are pulverized with a mortar and pestle and added to cold lysis buffer (0.01%Triton X100, 1x proteinase and phosphatase inhibitor cocktail, 1mM GTP in 1x BRB80 buffer) and left on ice for 20 minutes. Proteins are collected in the supernatant after centrifugation for 20 min at 15000 rpm. Tubulin (Cytoskeleton, Inc) is diluted to 10mg/ml in lysis buffer and incubated at 37 °C for 30 minutes, 100 μM taxol was added afterwards. Equal amounts of proteins from different mouse brains are warmed up to 37 °C and incubated with polymerized microtubules at 37 °C for 1 hour. Samples are centrifuged at 100,000x g at 37 °C for 40 minutes. Supernatants are saved and pellets are re-suspended in lysis buffer of the same volume of supernatant.

### *In Utero* Electroporation

*In utero* electroporation-mediated gene transfer was performed as previously described (Saito and Nakatsuji 2001; Tabata and Nakajima 2001). Briefly, E14.5 pregnant mice were anesthetized with ketamine/xylazine (100/10 mg/kg) and their uterine horn exposed. DNA plasmid (2-5 mg/ml) was injected via a pulled glass pipette into the lateral ventricle of each embryo, followed by electrodes placed on each side of the head parallel to the sagittal plane. Electrical current (five 50 ms pulses of 41 V with 950 ms intervals) was used to drive the plasmid DNA into lateral cortical areas. After sacrifice, mice were screened through visualizing of GFP expression using a stereo fluorescence microscope. GFP expressing mouse brains are dissected out and fixed in 3.7% paraformaldehyde for 3 hours. Samples are then transferred to PBS buffer with 30% sucrose and left at 4°C overnight. The mouse brains are sectioned at 20 μm using a Microtome (MICROM HM525).

### Western Analysis

Standard Western Blot analysis was performed using antibodies, as detailed above. The dual channel signal detection Licor system from Odyssey was used to analyze levels over a linear dynamic range.

### Constructs for protein expression in *E. coli*

The plasmids for 6His-PreScission-DCX-EGFP-StrepII (AddGene #83918), kif5b(1-560)-EGFP-6His (AddGene #15219), and Sfp-6His (AddGene #75015) were ordered from AddGene. The plasmid for JIP3 was a gift from Cavalli lab (Valeria Cavalli, Department of Anatomy and Neurobiology, Washington University in St Louis, School of Medicine, St Louis, MO, USA) (Sun et al., 2011). For DCX, EGFP was replaced by a ybbR-tag using Q5 mutagenesis (NEB #). For kif5b, the sequence encoding amino acids 1-490 was amplified with NdeI and EcoRI overhangs and inserted into a modified backbone based on pSNAP-tag(T7)2 (NEB #N9181S) before a SNAPf-EGFP-6His tag (Budaitis et al., 2021). For JIP3, the sequence encoding amino acids 3-240 (or 3-548) was amplified with NdeI-6His and EcoRI overhangs. The first two amino acids are Met in JIP3, which were therefore skipped because a 6His-tag was inserted at the N-terminus. The amplified sequence was then inserted into a modified backbone based on pSNAP-tag(T7)2 before a HaloTag-StrepII tag. All constructs were verified by restriction enzyme digestion and DNA sequencing.

### Protein expression in *E. coli*

Protein expression in *E. coli* was done as previously described (Budaitis et al., 2021). Briefly, a plasmid was transformed into BL21-CodonPlus(DE3)-RIPL competent cells (Agilent #230280), and a single colony was inoculated in 1 mL of TB with 50 µg/mL of chloroamphenicol and 25 µg/mL of carbenicillin or 15 µg/mL of kanamycin in the case of Sfp. The culture was shaken at 37°C overnight, and then inoculated into 400 mL of TB, which was shaken at 37°C for 5 hours, and subsequently cooled down to 18°C. IPTG was added to the culture to a final concentration of 0.1 mM, and the expression was induced overnight at 18°C with shaking. The culture was harvested by centrifugation at 3000 rcf for 10 minutes. Following the removal of the supernatant, the cell pellet was resuspended in 5 mL of B-PER complete (ThermoScientific #89821) supplemented with 4 mM MgCl2, 2 mM EGTA, 0.2 mM ATP, 2 mM DTT, and 2 mM PMSF. The cell suspension was then flash frozen and stored at –80°C.

### Purification and labeling of Sfp, JIP3, DCX, and KIF5B

The purification of *E. coli* expressed protein was done as previously described (Budaitis et al., 2021) . For the JIP3 and DCX constructs, a two-step purification was performed. For Sfp and kif5b, only the His-tag purification was performed. Briefly, the cell pellet was thawed at 37°C, and then nutated at room temperature for 20 minutes. The lysate was dounced for 10 strokes on ice and cleared via centrifugation at 80K rpm for 10 minutes at 4°C using a TLA 110 rotor (Beckman) in a tabletop Beckman ultracentrifuge. At the same time, 500 µL of the Ni-NTA slurry (Roche cOmplete His-Tag purification resin) was washed with 2x1 mL of wash buffer (WB, 50 mM HEPES, 300 mM KCl, 2 mM MgCl2, 1 mM EGTA, 1 mM DTT, 0.1 mM ATP, 1 mM PMSF, 0.1% (w/v) Pluronic F-127, pH 7.2) in a 10-mL column (Bio-Rad #7311550). After the centrifugation, the supernatant was loaded into the column, and allowed to flow through the resin by gravity. The resin was then washed with 3x2 mL of WB.

For Halo-tag labeling, halo-tag ligand was added to final 10 µM, and the resin was incubated at room temperature for 10 minutes. For ybbR-tag labeling, the CoA-dye ligand for ybbR-tag was generated by reacting Coenzyme A (CoA) with a dye containing a maleimide group in 1:1 ratio at room temperature for 30 minutes. The final product was quenched with 50 mM DTT, aliquoted, flash frozen, and stored at –80°C. To label the ybbR-tag, CoA-dye and Sfp was added to the resin to a final concentration of 10 µM. The resin was nutated at 4°C for 3 hours. After the labeling, the resin was washed with 3x3 mL of WB, and eluted with Ni-NTA elution buffer (Ni-EB, 50 mM HEPES, 150 mM KCl, 2 mM MgCl2, 1 mM EGTA, 1 mM DTT, 0.1 mM ATP, 1 mM PMSF, 0.1% (w/v) Pluronic F-127, 250 mM imidazole, pH 7.2). For BirA and kif5b, the protein was aliquoted, flash frozen, and stored at –80°C until further usage. For JIP3 and DCX, the concentrated fraction was pooled and flown through 1 mL of streptactin slurry (IBA #2-1201) which had been washed with 2x1 mL WB. The resin was then washed with 3x2 mL WB, and then eluted with streptactin elution buffer (St-EB, 50 mM HEPES, 150 mM KCl, 2 mM MgCl2, 1 mM EGTA, 1 mM DTT, 0.1 mM ATP, 1 mM PMSF, 0.1% (w/v) Pluronic F-127, 2.5 mM dethiobiotin, pH 7.2). The concentrated fraction was pooled and further concentrated via centrifugation using Amicon 0.5-mL 10 kDa unit. The protein was verified on a PAGE gel, and the concentration was determined using Braford assay.

### Constructs for protein expression in insect cells

The pFastBac plasmid with codon-optimized full-length human dynein was a gift from the Carter lab (MRC laboratory of Molecular Biology, Francis Crick Avenue, Cambridge, UK) (Schlager et al, 2014). The pFastBac plasmid that encodes tail-truncated human dynein (amino acids 1320-4646 of DYNC1H1) was a gift from the Reck-Peterson Lab (Department of Cellular and Molecular Medicine, University of California, San Diego, CA, US) (Htet et al, 2020).

### Protein expression in insect cells

Full-length human dynein and tail-truncated human dynein were expressed in Sf9 cells as described previously (Schlager et al, 2014; Htet et al, 2020). Briefly, the pFastBac plasmid containing full-length human dynein or tail-truncated dynein was transformed into DH10Bac competent cells (Gibco, #10361012) with heat shock at 42 °C for 45 seconds followed by incubation at 37 °C for 4 hours in S.O.C. medium (Gibco, #15544034). The cells were then plated onto LB agar plates containing kanamycin (50 μg mL^−1^), gentamicin (7 μg ml^−1^), tetracycline (10 μg mL^−1^), BluoGal (100 μg mL^−1^) and isopropyl-β-D-thiogalactoside (IPTG; 40 μg mL^−1^), and positive clones were identified by a blue/white color screen after 36 hours. Bacmid DNA was extracted from overnight culture using an isopropanol precipitation method with Qiagen buffer (Qiagen, #27104) as described previously (Schlager et al, 2014). To generate baculovirus for Sf9 insect cell transfection, 2 mL of Sf9 cells at 0.5×10 ^6^ cells per mL in six well plates (Corning, #3516) were transfected with 2 μg of fresh bacmid DNA and 6 μL of FuGene HD transfection reagent (Promega, E2311) according to the manufacture’s instruction. The cells were incubated for 4 days and the supernatant containing V0 virus was collected then. To generate V1 virus, 0.5 mL of V0 virus was used to transfect 50 mL of Sf9 cells at 1.5 × 10^6^ cells per mL. The supernatant containing V1 virus was collected by centrifugation at 200 g for 5 minutes at 4 °C after 3 days. The V1 virus was stored at 4 °C in the dark until use. For protein expression, 5 mL of the V1 virus was used to transfect 500 mL Sf9 cells at 2 × 10^6^ cells per mL. After 60 hours incubation, cells were collected by centrifugation at 3000 g for 10 minutes at 4 °C. The cell pellet was resuspended in 15 mL ice -cold PBS and centrifuged again. The supernatant was then removed, and the cell pellet was flash-frozen in liquid nitrogen and stored at -80 °C.

### Purification and labeling of tail-truncated and full-length human dynein

Full-length dynein and tail-truncated dynein was purified from frozen Sf9 pellets as described previously (Schlager et al, 2014; Htet et al, 2020). Frozen pellets from 500 mL insect cell culture were thawed on ice and resuspended in lysis buffer (50 mM HEPES pH 7.4, 100 mM NaCl, 1 mM DTT, 0.1 mM ATP, 10% (v/v) glycerol, 2 mM PMSF) supplemented with 1 protease inhibitor cocktail tablet (cOmplete-EDTA free, Roche, #11836170001) to a final volume of 50 mL. Cells were then lysed using a Dounce homogenizer with 20 strokes. The lysate was cleared by centrifugation at 279,288 g for 10 minutes at 4 °C using a Beckman Coulter tabletop centrifuge unit. The clarified supernatant was incubated with 3 mL of IgG Sepharose 6 Fast Flow beads (Cytiva, #17096901) for 4 hours with rotation. After incubation, the protein bound IgG beads were transferred to a gravity flow column and washed with 100 mL lysis buffer and 100 mL TEV buffer (50 mM Tris-HCl pH 8.0, 250 mM potassium acetate, 2 mM magnesium acetate, 1 mM EGTA, 1 mM DTT, 0.1 mM Mg-ATP and 10% (v/v) glycerol). To fluorescently label the carboxy-terminal SNAPf tag of full-length human dynein, dynein coated beads were incubated with 5 μM SNAP-Cell TMR (New England BioLabs, #S9105S) in the column for 10 minutes at room temperature. The beads were then washed with 100 mL TEV buffer at 4 °C to remove unbound dyes. Subsequently, the beads were resuspended in TEV buffer (final volume 5 mL) with 100 μL TEV protease (New England BioLabs, #P8112S) and incubated at 4°C on a roller overnight. After TEV cleavage, the beads were removed and protein of interest was concentrated using a 100 kDa molecular weight cut-off (MWCO) concentrator (Millipore, #Z648043) to 1 mL and flash-frozen in liquid nitrogen.

### Microtubule polymerization

Microtubule polymerization was performed as described before (Rao et al., 2018). Briefly, 2 µL of 10 mg/mL unlabeled tubulin (Cytoskeleton) was mixed with 2 µL o f 1 mg/mL biotin-tubulin and 2 µL of 1 mg/mL dye-labeled tubulin on ice. 0.5 µL of 10 mM GTP was added to the mixture, and the mixture was incubated at 37°C for 20 minutes. Afterwards 0.7 µL of 0.2 mM taxol (in DMSO) was added, and the solution was incubated at 37°C for another 15 minutes. The un -incorporated tubulin was removed by centrifuging through a glycerol cushion (80 mM PIPES, 2 mM MgCl2, 1 mM EGTA, 60% glycerol, 10 µM taxol, 1 mM DTT, pH 6.8) at 80k rpm for 5 minutes at room temperature using TLA motor in a tabletop Beckman ultracentrifuge. The supernatant was discarded, and the pellet was resuspended in 12 µL resuspension buffer (80 mM PIPES, 2 mM MgCl 2, 1 mM EGTA, 10% glycerol, 10 µM taxol, 1 mM DTT, pH 6.8) to obtain a final 2 mg/mL MT concentration for the TIRF assay. For the MT-binding and -release assay, 5 µL of 10 mg/mL unlabeled tubulin was used to polymerize the MTs, and the pellet was resuspended in 10 µL of the resuspension buffer to obtain a final 5 mg/mL MT concentration.

### Microtubule-binding and -release assay of kif5b and single-head human dynein

Impaired/inactive motors were removed by a MT-binding and -release assay as described before (Budaitis et al., 2021; Rao et al., 2019). Briefly, 50 µL of protein solution was exchanged into a binding buffer (30 mM HEPES, 50 mM KCl, 2 mM MgCl2, 1 mM EGTA, 10% glycerol, 1 mM DTT, 0.1 mM AMP-PnP, pH 7.2) using Zeba 7-kDa unit (ThermoScientific #89882). The protein solution was warmed to room temperature, and taxol was added to final 20 µM concentration. For kif5b, AMP-PnP was also added to a final 1 mM concentration. 3 µL of the 5 mg/mL MT stock was added to the protein solution and then mixed well. The solution was then carefully layered on top of 100 µL of glycerol cushion for kif5b (80 mM PIPES, 2 mM MgCl2, 1 mM EGTA, 60% glycerol, 10 µM taxol, 1 mM DTT, pH 6.8) or sucrose cushion for dynein (30 mM HEPES, 50 mM KCl, 2 mM MgCl2, 10% glycerol, 25% w/v sucrose, 10 µM taxol, 1 mM DTT, pH 7.2) in a TLA 100 rotor (Beckman), and centrifuged at 45krpm for kif5b or 80krpm for dynein at room temperature for 10 minutes. Afterwards, the supernatant was removed and the pellet was washed with 2x20 µL wash buffer (30 mM HEPES, 50 mM KCl, 2 mM MgCl2, 10% glycerol, 10 µM taxol, 1 mM DTT, pH 7.2). The pellet was then resuspended in 47 µL of high-salt release buffer (HSRB, 30 mM HEPES, 300 mM KCl, 2 mM MgCl2, 10% glycerol, 10 µM taxol, 1 mM DTT, pH 7.2), and 3 µL of 100 mM ATP was added to the solution. The solution was centrifuged at 40k rpm for 5 minutes, and the supernatant was aliquoted, flash frozen, and stored at –80°C for further usage.

### DDJ complex assembly

DDJ complex was assembled following a published protocol (Potokar et al.). Briefly, dynein, dynactin (a gift from the laboratory of Andrew Carter, MRC), and JIP3 were mixed on ice in 1:1:1 ratio (final concentration of 200 nM each) and incubated for 1 hour on ice in the dark. For DDJ complex formation in the presence of N-DCX, N-DCX was added in equal amount as dynein.

### TIRF motility assay

The TIRF motility assay was performed as described before (Budaitis et al., 2021). Briefly, a coverslip was cleaned using ethanol, and assembled into a flow chamber. 10 µL of 0.5 mg/mL biotin-BSA was introduced into the flow chamber, and the flow chamber was incubated at room temperature for 10 minutes in a humidity chamber. The chamber was then washed with 3×20 µL of blocking buffer (BB, 80 mM PIPES, 2 mM MgCl 2, 1 mM EGTA, 10 µM taxol, 1% (w/v) Pluronic F-127, 2 mg/ml BSA, 1 mg/mL α-casein, pH 6.8), and incubated for 10 minutes. The solution in the chamber was completely removed using vacuum, and 10 µL of 0. 25 mg/mL streptavidin was flown in and incubated at room temperature for 10 minutes. The chamber was then wash with 3×20 µL of BB, and the solution was completely removed afterward. 0.5 µL of 0.2 mg/mL fluorescently labeled MTs was diluted in 19.5 µL of BB and flown into the chamber. The chamber was then washed with 2x20 µL BB and 20 µL of motility buffer (MB, 60 mM HEPES, 50 mM KAc, 2 mM MgCl2, 1 mM EGTA, 0.5% (w/v) Pluronic F-127, 10 µM taxol, 1 mM DTT, 5 mg/mL BSA, 1 mg/mL α-casein, pH 7.2). 1 µL of 100 mM ATP, 1 µL of 50 mM biotin, and 1 µL of oxygen scavenger system was added to 46 µL of MB, and 1 µL of 10 nM DDJ complex was added subsequently. For DCX and N-DCX experiments, DCX was added to a final 10 nM in the final solution. The solution was mixed well and flown into the chamber. The chamber was then sealed with vacuum grease. The acquisition time was 200 ms per frame, and a total of 600 frames was taken for each movie. The data was analyzed using a custom-built MATLAB software, and the statistical analysis and data visualization were performed using Prism.

### TIRF gliding assay

A slide chamber was assembled as described above. 10 µL of 0.1 mg/mL anti -GFP antibody (YenZym) was introduced into the chamber, which was then incubated in a humidity chamber for 10 minutes. The chamber was washed with 3×20 µL BB and 20 µL of MB. 1 µL of MTBR fraction of single-head human dynein was diluted in 19 µL of MB, and the solution was flown into the chamber and incubated for 2 minutes. The chamber was washed with 3×20 µL of MB to remove unbound dynein. 1 µL of 100 mM ATP, 0.5 µL of 0.2 mg/mL MTs, and 1 µL of oxygen scavenger system was added to 47.5 µL of MB, which was flown into the chamber. The chamber was sealed with vacuum grease. The imaging condition and analysis was done as described above.

### Statistical Analysis

Statistical analyses were performed using GraphPad Prism 7.0 software (GraphPad Software Inc., San Diego, CA, USA). All data are presented as the mean ± SEM of at least three independent experiments. Statistical significance was determined using one-way analysis of variance (ANOVA) followed by Tukey’s test if more than two groups were analyzed. Two-tailed test and student’s t-test was used to compare two groups. P < 0.05 was considered significant (* p < 0.05; # p < 0.01, if not specified otherwise).

**Supplemental video1.** Live-cell imaging shows DIC mobility in a WT neuron.’

**Supplemental video2.** Live-cell imaging shows DIC mobility in a Dcx-/y neuron.

**Supplemental video3.** Live-cell imaging shows TrkB mobility in a WT neuron.

**Supplemental video4.** Live-cell imaging shows TrkB mobility in a Dcx-/y neuron.

**Supplemental video5.** Live-cell imaging shows TrkB mobility in a Dcx-/y neuron transfected with JIP3-shRNA.

**Supplemental video6.** Live-cell imaging shows TrkB mobility in a WT neuron transfected with JIP3.

**Supplemental video7.** JIP3 has a transient affinity for dynein.

## AUTHOR CONTRIBUTIONS

X.F., J.S.L., L.R., and A. G. conceived and designed this study, assisted with data analysis and interpretation, and wrote the manuscript. X.F. and P.L. performed time-lapse imaging, immunoprecipitation, western analysis, pull-down assays, and related data analysis. Q.W. performed in pull-down assays. In utero electroporations were performed and analyzed by P.L., X.F., and A.S.. L.R. generated and purified JIP3 and DCX-ybbR constructs, and performed the *in vitro* TIRF motility assays and related data analyses. X. L. expressed and purified tail-truncated and full-length human dynein.

## ACKNOWLEDGEMENTS

The authors thank Lisa Baker and Julian Curiel for help with the editing the manuscript. L.R. and A.G thank Andrew Carter (LMB Cambridge) for generously providing purified porcine brain dynactin. This work was supported by the Brain and Behavior Research Foundation (J.S.L. and M.T.), the Whitehall Foundation (J.S.L.), the National Natural Science Foundation of China (81971425 and 81871035) (X.F. and P.L.), the Zhejiang Provincial Natural Science Foundation of China (LZ09H090001 and LY20H040002) (P.L. and X.F.), the National Institute of Health (NIH) grants R01GM098469 and R01NS114636 (L.R, X.L. and A.G.), and the NIH grant RO1NS104428-01 (J.S.L.).

---

**Supplemental Figure 1.**
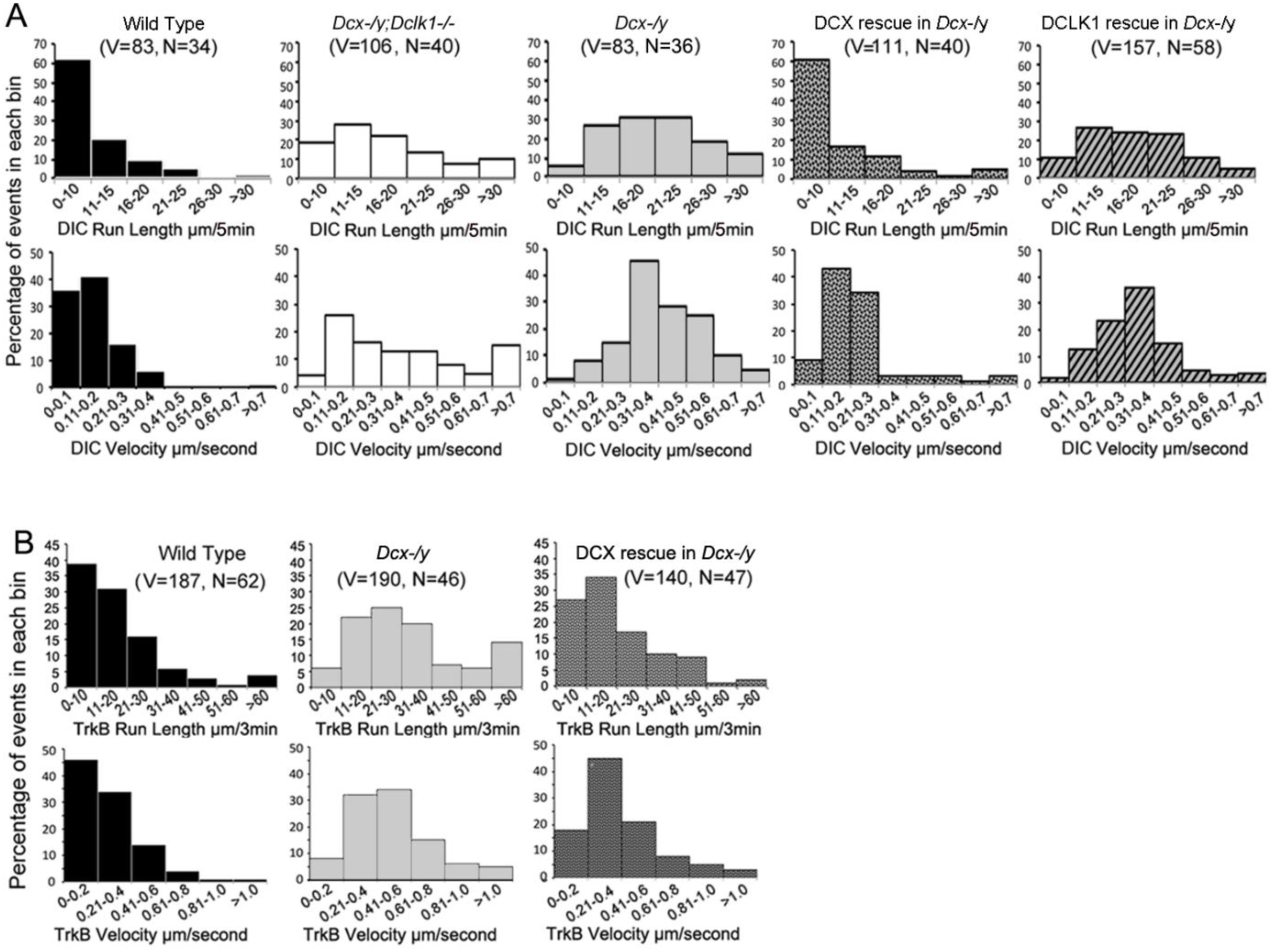
Run length and velocity distributions of DIC and TrkB in different neurons. (A) Run length and velocity distributions of retrograde DIC complexes in different neurons are shown. Total numbers of neurons (N) and vesicles (V) used in the calculations are indicated in the panel. (B) Run length and velocity distributions of retrograde TrkB complexes in different neurons are shown. Total numbers of neurons (N) and vesicles (V) used in the calculations are indicated in the panel.

**Supplemental Figure 2.**
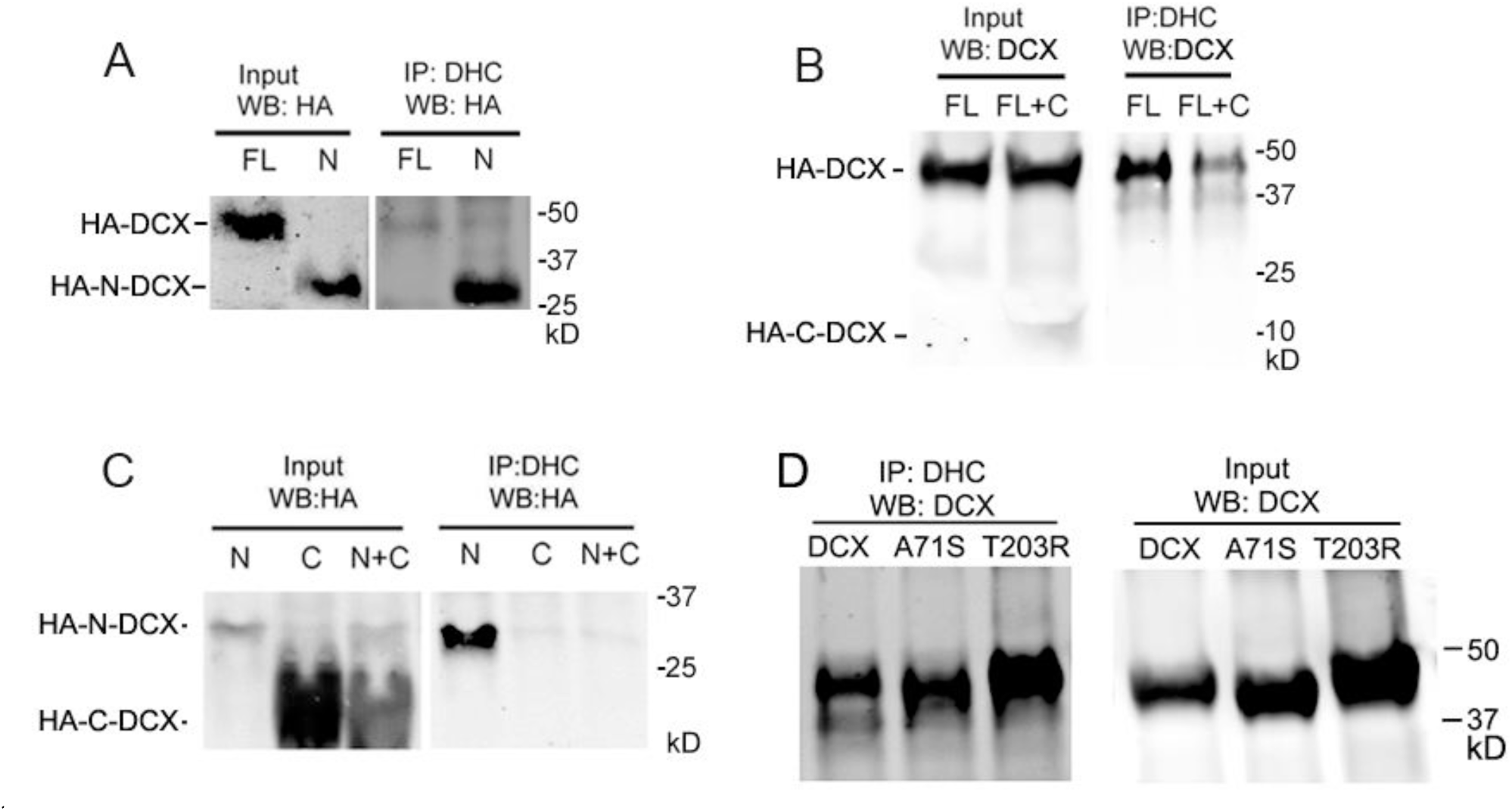
DCX associates with dynein heavy chain through its N-terminal domain. (A) HEK cells are transfected with full-length DCX (FL) or N-DCX (N) for two days. Protein lysates were used for immunoprecipitation and analyzed by Western blot as indicated in the figure. More N- DCX proteins are precipitated with DHC compared to full-length DCX, while similar amount of full-length DCX and N-DCX is expressed in transfected cells. (B) HEK cells are transfected with full-length DCX (FL) with/without C-DCX (C) for two days. The presence of C-DCX decreases association of full-length DCX with DHC. (C) Similarly, C-DCX decreases association of N-DCX with DHC. C-DCX does not bind DHC.

**Supplemental Figure 3.**
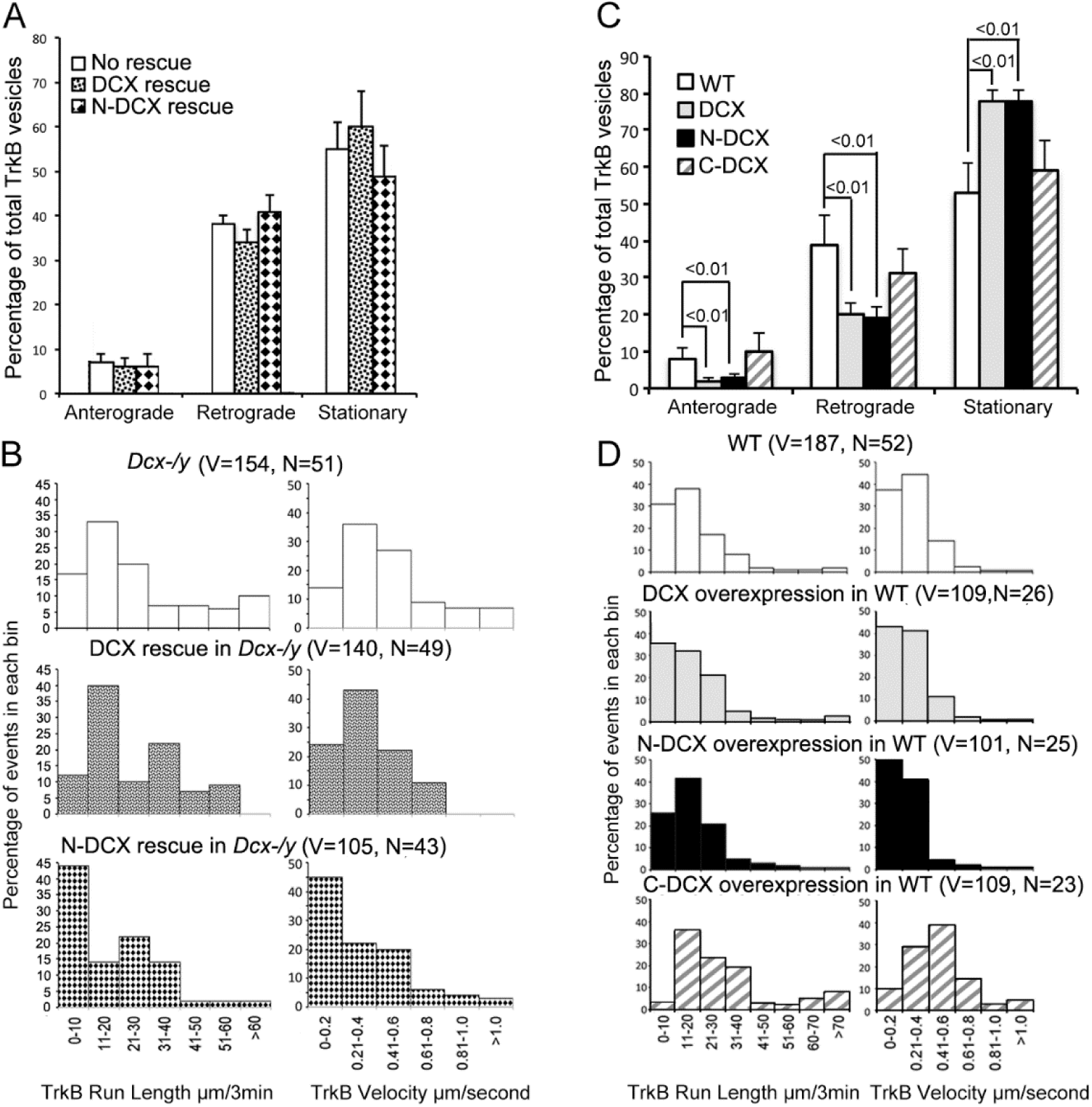
DCX affects the retrograde transport through DCX/dynein interaction. (A) Distribution calculations of the TrkB vesicle mobility status (anterograde, retrograde, and stationary) are demonstrated. No significant differences are observed among different neurons. (B) Run length and velocity distributions of retrograde TrkB complexes in axons from different neurons are shown. Total numbers of neurons (N) and vesicles (V) used in the calculations are indicated in the panel. (C) Distribution calculations of the TrkB vesicle mobility status (anterograde, retrograde, and stationary) in axons from WT cells with different transfections are demonstrated. Overexpression of DCX or N-DCX significantly increased percentage of stationary TrkB vesicles in axons, while decreased the percentile of both anterograde and retrograde transport. (D) Run length and velocity distributions of retrograde TrkB complexes in axons from different neurons are shown. Total numbers of neurons (N) and vesicles (V) used in the calculations are indicated in the panel.

**Supplemental Figure 4.**
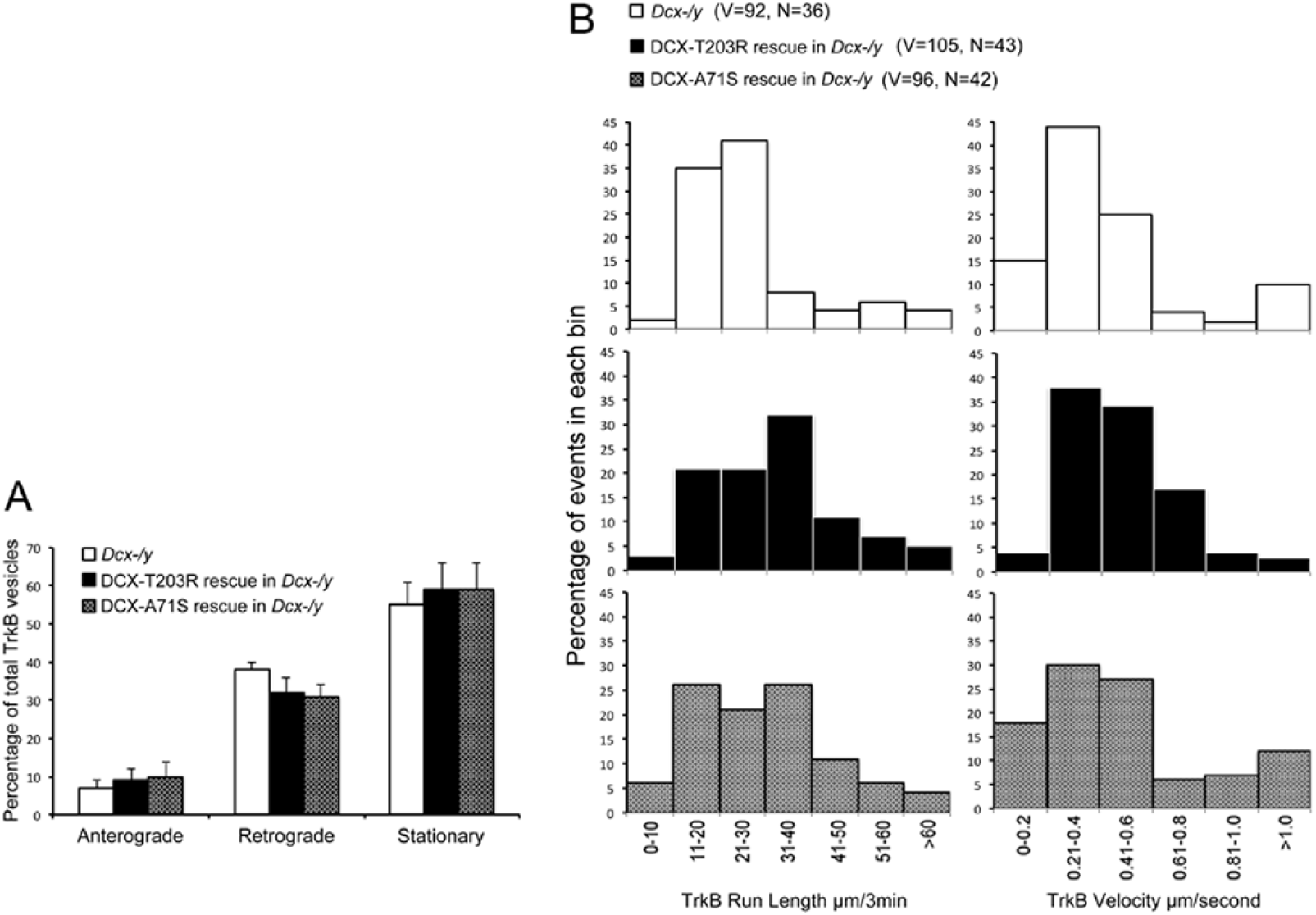
DCX effect on the retrograde trafficking needs DCX/MT interaction. (A) Distribution calculations of the TrkB vesicle mobility status (anterograde, retrograde, and stationary) in different cells are demonstrated. No significant differences are observed among different neurons. (B) Run length and velocity distributions of retrograde TrkB complexes in axons from different neurons are shown. Total numbers of neurons (N) and vesicles (V) used in the calculations are indicated in the panel.

**Supplemental Figure 5.**
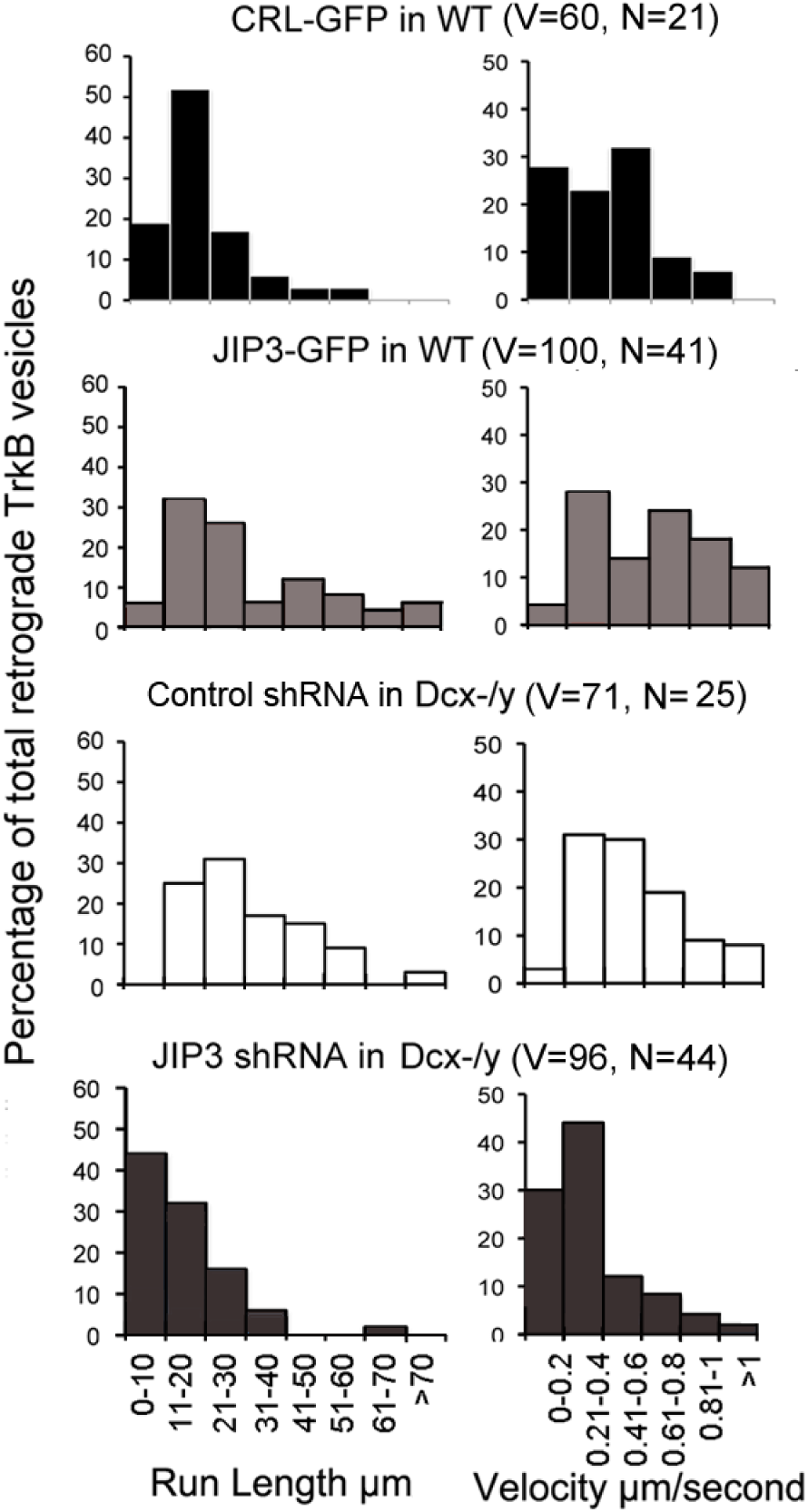
JIP3 enhances retrograde transport of TrkB. Run length and velocity distributions of retrograde TrkB complexes in axons from different neurons are shown.’

**Supplemental Figure 6:**
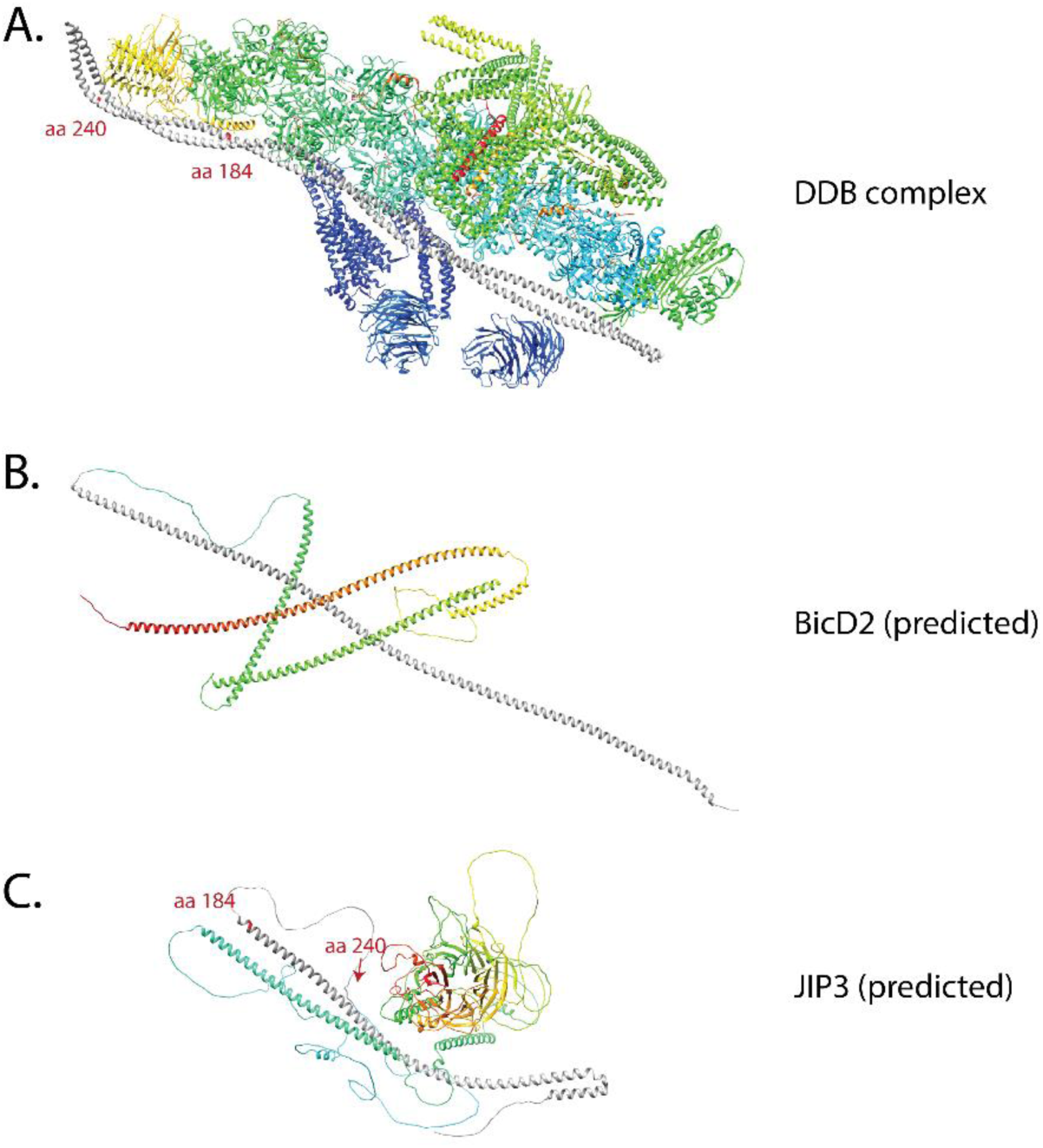
Comparison of the predicted structures of BicD2 and JIP3 with cryo-EM structure of DDB containing BicD2. (A) Cryo-EM structure of DDB complex (PDB 5AFU) (Urnavicius et al., 2015). The structure shows that the first coiled coil of BicD2 (dark gray) is up to aa 275, which spans the full length of dynactin shoulder. (B) Predicted BicD2 (Uniprot Q8TD16) structure. The dark gray indicated the first α-helix that was predicted to extend to aa 272 (Jumper et al., 2021), which corresponds to the cryo-EM structure. (C) Predicted JIP3 (Uniprot Q9UPT6) structure. The dark gray indicated the first α-helix is predicted to be up to aa184. The region between aa 185 and aa 240 was predicted to be disordered. The red labels in (A) indicate the beginning (aa 6) of the BicD2 α-helix, the position of aa 187 to show the estimated end of the predicted α-helix in JIP3 in (C), and the position of aa 240 to show the estimated aa 240 position of JIP3, should the disorder region of JIP3 forms α-helix upon interaction with dynactin. The red labels in (B) and (C) indicates the beginning and end of the predicted first α-helix in BicD2 and JIP3, respectively.

**Supplemental Figure 7:**
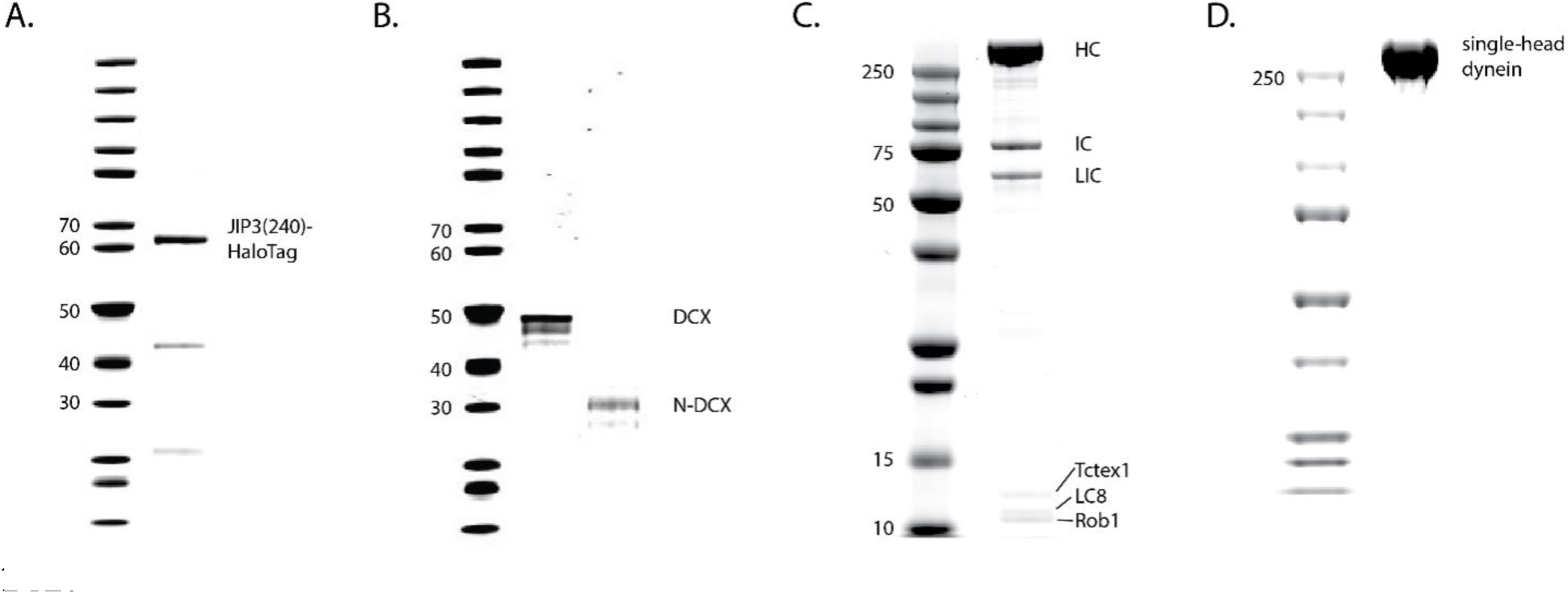
PAGE gels of recombinant expressed proteins. (A) JIP3(aa1-240)-HaloTag. (B) DCX-ybbR and N-DCX-ybbR. (C) Human dynein complex, containing a heavy chain (HC) with a SNAP-tag, an intermediate chain (IC), a light intermediate chain (LIC), and three light chains (Tctex1, LC8, Rob1). (D) Single-head human dynein-GFP.

**Supplemental Figure 8:**
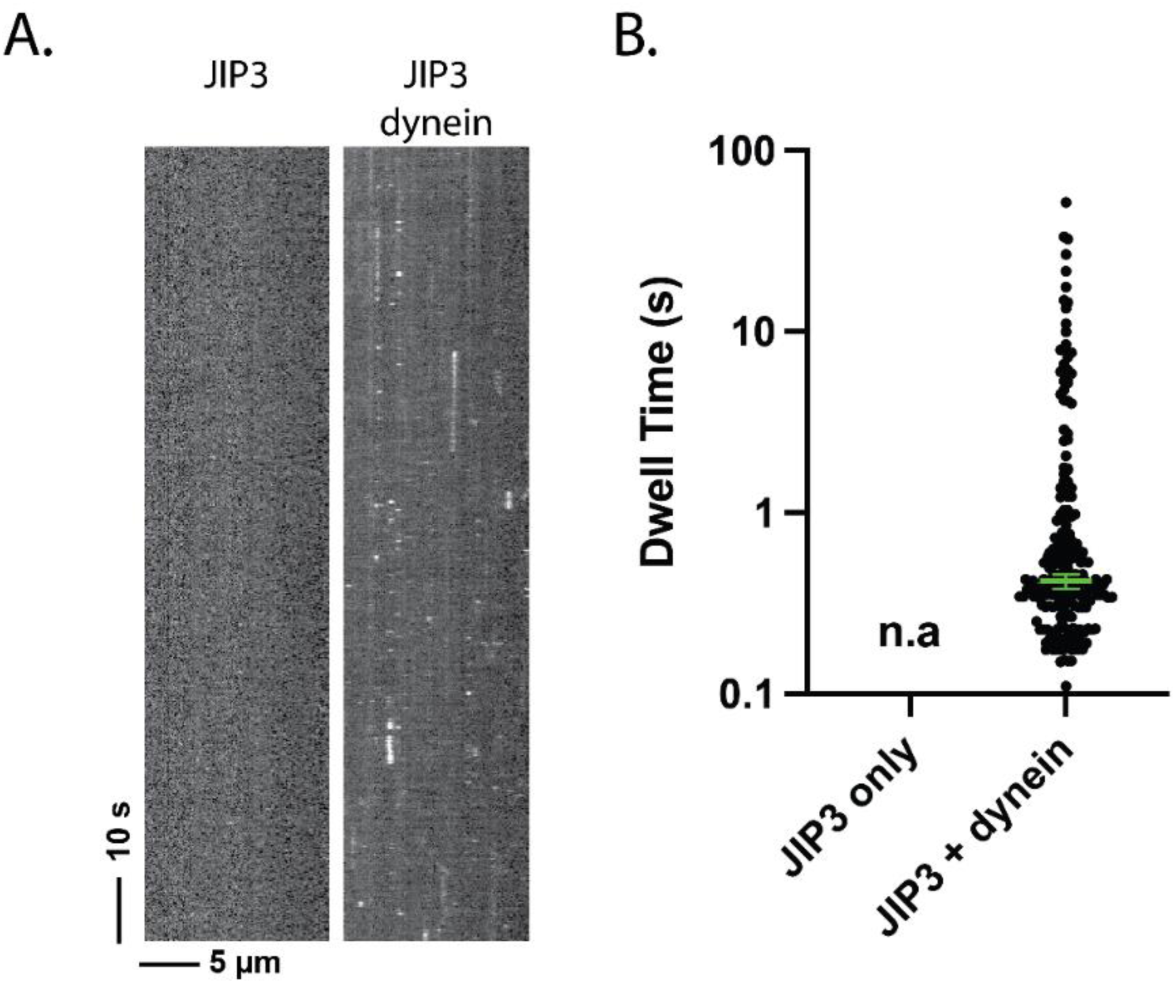
JIP3 has a transient affinity for dynein. (A) Kymograph of JIP3 on microtubules, without (left) or with (right) dynein present. Without dynein, JIP3 shows no affinity for microtubules; in the presence of dynein, JIP3 demonstrated brief binding events via dynein. The concentration of dynein was 1 nM, and the concentration of JIP3 was 10 nM. (B) The dwell time JIP3 on microtubule via dynein. Without dynein, there was no measurable dwelling of JIP3; with dynein, the dwell time of JIP3 on dynein is 0.42 [0.38, 0.46] s (median [95% CI]).

**Supplemental Figure 9.**
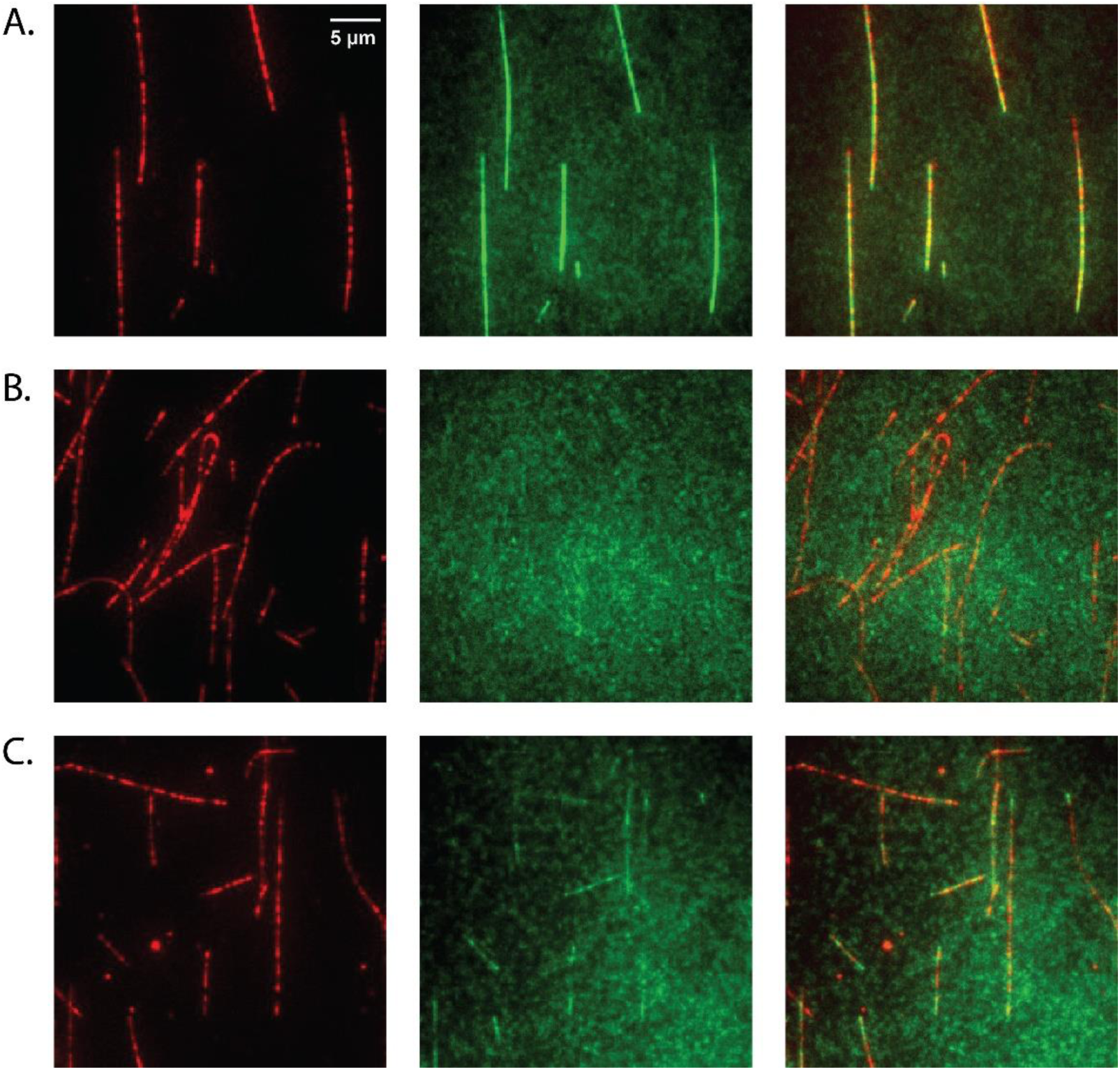
DCX binds MTs. (A) At 10 nM concentration, DCX uniformly decorates microtubules. DCX-ybbR was labeled with CoA-CF488, and microtubules were labeled with Cy5. (B) At 10 nM concentration, N-DCX doesn’t bind to microtubules in the motility buffer. N-DCX-ybbR was labeled with CoA-CF488. (C) At 10 nM concentration, N-DCX decorates microtubule in a buffer that is half of the ionic strength of motility buffer.

